# Enviromic prediction is useful to define the limits of climate adaptation: A case study of common beans in Brazil

**DOI:** 10.1101/2022.02.07.478581

**Authors:** Alexandre Bryan Heinemann, David Henriques da Matta, Igor Kuivjogi Fernandes, Roberto Fritsche-Neto, Germano Costa-Neto

## Abstract

Future environmental shifts foster plant research aiming to develop climate-smart cultivars but the past and current impacts on the environment are also a key to unraveling a major part of the phenotypic adaptation of crops. These studies may determine the most relevant environmental components of yield stability and adaptability within a breeding framework. Here, as a proof-concept study, we quantified the impacts of climate drivers in adapting common bean across Brazilian regions and seasons. We developed an ‘enviromic prediction’ approach based on Generalized Additive Models (GAM), large-scale environmental covariate data (EC), and grain yield (GY) of 18 years of a common bean breeding program. Then, we predicted the optimum limits for ECs for each production scenario. We verified the ability of GAM-based models to explain the climate driver GY variation and performed accurate predictions for diverse production scenarios (four regions, three seasons, and two grain types). Our results indicates that air temperature (maximum and minimum), accumulated solar radiation, and rainfalls are mostly associated as the main drivers of GY variation in most regions. We also observed a huge variability of the climate drivers impact for the same germplasm cultivated across different seasons for each region. Furthermore, this climate influence in common beans adaptation is more evident during the vegetative for some seasons, while more impressive for reproductive stages for other seasons. Consequently, it demands higher efforts from breeding programs in developing region- or season-specific ideotype cultivars. Enviromics prediction with GAM was useful to identify the effect of climate on critical crop stages, which indirectly might help breeders in developing climate-smart varieties. We envisage its use with research field trial data (e.g., advanced yield testing) and historical farm field yield aimed at understanding breeding gaps in developing adapted cultivars for growing scenarios.

**Highlights:**

- We developed an ‘enviromic prediction’ approach based on Generalized Additive Models (GAM) for a large-scale environmental covariate data and grain yield
- We verified the ability of GAM-based models to explain the climate driver grain yield variation and performed accurate predictions for diverse production scenarios (four regions, three seasons, and two grain types)
- Climatic limitations for cropping commun beans were identified across seasons and regions.
- “Optimum” values for climate variables in different common bean regions productions were obtained .

## 1 INTRODUCTION

Legumes contain almost two to three times more proteins than cereals. Their dietary importance as a protein source is well documented. Among 1,300 species of legumes, common beans (Phaseolus vulgaris L.) is the most consumed worldwide (Sathe, 2016). Brazil ranks third in terms of legume production, with 3.126 million metric tons of grains expected for the 2020/2021 crop season (CONAB, 2020), and ranks second in terms of harvested area (ca. 2.9 million ha planted). Its consumption was ca. 15 kilograms per person in 2019. This production basically comprises two main types of common beans (Phaseolus vulgaris L): the “Carioca” and “Black” types, corresponding to 70% and 20% of the total production and representing 61% and 17% of the Brazilian consumption, respectively (Silva, 2019; Souza et al., 2020).

Although abiotic pressures on productivity are difficult to calculate accurately, abiotic stresses exert a significant effect on agricultural production depending on the class of damage to the total cultivated area (Al-Tawaha, 20201). The major limiting factors in common bean production are drought stress, low soil fertility (Beebe et al., 2011; Müller et al., 2014), and nitrogen deficiency due to poor nitrogen fixation (Rao, 2001). In the future, a general trend towards temperature homogenization will act to homogenize common bean yields and increase drought frequency (Heinemann et al., 2017).

In Brazil, the common bean production extends from the South to the North and spreads across three distinct growing seasons: wet, dry, and winter. Wet and dry growing seasons are rainfed systems and represent 93% of the common bean Brazilian production area, whereas the winter sowing is fully irrigated (Heinemann et al., 2016). Due to environmental variability, the performance of cultivars varies substantially across regions and seasons. Maximizing genetic gains in producing regions requires a better environment characterization aiming to produce information that may assist breeding strategies in developing adapted yield germplasm for a producing region.

Despite recommendations that go back to the 1960s, the use of environmental information in predictive and in exploratory models is little used, and most plant breeders (Cooper et al., 2014; Xu, 2016) prefer indirect measurements of environmental quality (e.g., use of average values for all genotypes for a certain trait in a certain environment; Finlay and Wilkinson, 1963). The first models considering environmental information involved simple linear regression models (e.g., FR, Wood, 1976; Denis, 1980; Romay et al., 2010), partial least squares (e.g., PLS, Vargas et al., 1999; Monteverde et al., 2019; Poker et al., 2020), and bootstrap techniques with FR and geographic information tools (FR-GIS, Costa-Neto et al., 2020). A promising approach combining the aspects of predictive and exploratory models in a single and efficient manner is the Generalized Additive Model (GAM, Hastie; Tibshirani, 1986). This approach was developed as an alternative to the generalized linear model (GLM) (Nelder and Wedderburn, 1972) and allowed the improvement of non-parametric functions as possible predictors (Arnold et al., 2019).

The advance of enviromics technique (Resende et al., 2021; Costa-Neto et al., 2021a), mostly because of readily available field sensors and open-source pipelines (e.g., EnvRtype, Costa-Neto et al., 2021b), allows implementing expensive computational tools that deal with big data of envirotyping and phenotyping (Cooper et al., 2014; Crossa et al., 2021). The latter is one of the most important databases for plant research and relies on the historical field trial records of plant breeding. Using this extensive database allows going deeper into the impacts of the environment on phenotypic variation, which may guide plant breeders to dealing with unknown sources of G×E that limit the targeting of cultivars and genetic progress in some regions (Annicchiarico, 2002, Xu, 2016).

Here, we provide a broader assessment of how climate variables may help breeders to develop climate-smart varieties using as a case study the common bean yield in several production regions in Brazil. First, we introduce the concept of “climate prediction” as an extension of the so-called “enviromics prediction” (Resende et al., 2021; Costa-Neto et al., 2021b), here implemented using the GAM approach and involving large-scale phenotypic and environmental data. Then, we analyze the main climate factors in each region, season, and grain type of common bean in Brazil in order to train the most accurate prediction model and verify the impacts of the most important factors for common bean variation. Finally, we discuss climate-oriented limits of adaptation of common beans and the viability of the GAM-based model supplied with climate data in performing predictions of grain yield. We then discuss the benefits of this approach in using climate data in the decision-making process in testing, selecting, and targeting common bean cultivars at a national level in Brazil.

## 2 MATERIAL AND METHODS

### 2.1 Regional setting

The study area encompasses a wide range of locations, soils, and rainfall zones (Midwest, South, Southeast, and Northeast regions) cultivated with common beans (Fig 1A). The study areas include four biomes (“Cerrado,” “Caatinga,” “Mata Atlântica,” and “Pampa”). The altitude, latitude, and longitude are 0-1,200 m above mean sea level, 7° S to 29° S, and 36° W to 58° W (Figure 1B), respectively. The study area concentrates 98% of common dry bean production (IBGE, 2017). Basically, two types of common bean are cultivated, the so-called “Carioca” and “Black.”

**Figure 1.**
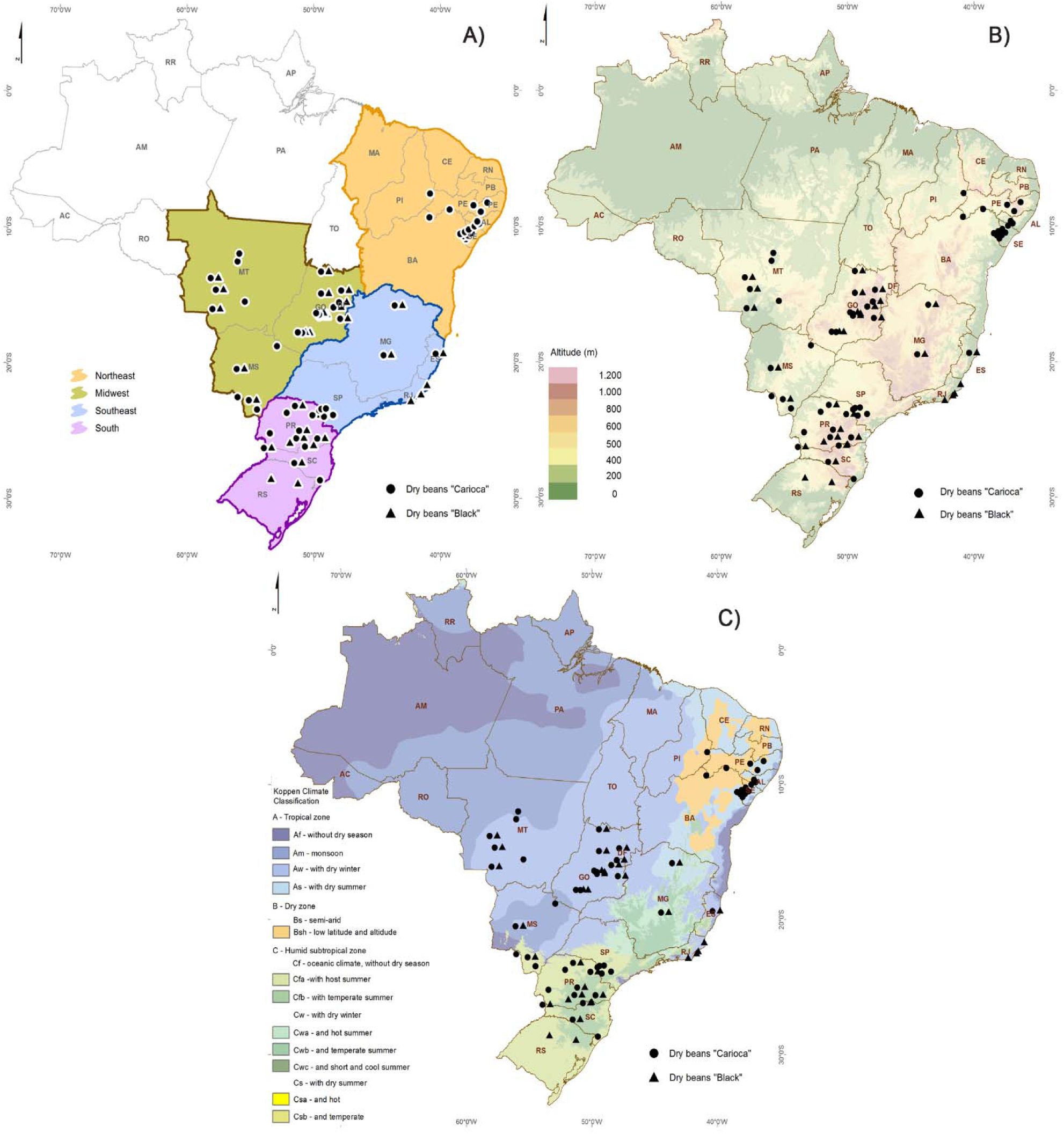
Geographic location of common bean trials in Brazil. **A**) South, Midwest, Southwest, and Northeast production regions; **B**) Altitude (m); and **C**) Köppen climate classification of Brazilian production regions. For all panels, black triangle dots and circle dots correspond to the “Black” and “Carioca” (circle) grain types, respectively.

The growing environments located in the Southeast (blue colors in Fig 1A) and South (purple colors in Fig 1A) regions are the most socially and economically important production sites, accounting for 28% and 27% of the national production, respectively. In those regions, the common bean production is mainly spread out in the humid tropical zone (Cfa and Cfb) (Fig 1C) and characterized by having two mainly common bean crop seasons. For the Southeast region, there are wet (accounting for 36% of the common bean production region) and winter (accounting for 33% of the production region) seasons, and for the South region, there are wet (46% of the production) and dry seasons (53% of the production). In the Southeast region, sowing dates are from August to September (dry season) and October to December (wet season), and for South region, sowing dates are from August to September (wet season) and October to November (dry season).

The Midwest region accounts for 22% of the Brazilian production. The predominant climate in this region is tropical wet and dry, or savannah (Aw), which represents 94%, 52.8%, and 36.6% of the total area of Goiás, Mato Grosso, and Mato Grosso do Sul States, respectively (Alvares et al., 2013). This region comprises three seasons of common beans: wet, dry, and winter, representing 21, 34, and 44% of the total common bean produced in the region (IBGE, 2017). Sowing for the wet, dry, and winter seasons occurs between the beginning of November and the end of December, the middle of January and February, and from May to June, respectively (Heinemann et al., 2017).

The Northeast region accounts for 20% of the Brazilian production. In this region, common bean production mainly concentrates in the tropical zone (As) (Figure 1C). This region is characterized by two common bean seasons, wet and winter, and they represent 67% and 33% of the total common bean produced in the region (IBGE, 2017). Sowing for wet and winter seasons occurs between October and December and from January to May. The main common bean type produced is “Carioca.”

### 2.2 Experimental Data Set

This study uses a large, accumulated yield data set (424 trials) formed by variety trials on commonly grown and well-adapted common bean varieties in Brazil from 2011 to 2018 (Figure 1). This set of variety trials were conducted as a multi-environmental (MET) condition as a part of the EMBRAPA common bean breeding program in Brazil. This MET for variety testing and verifying the performance of candidate varieties is divided into two common bean types, according to the grain type most consumed in Brazil: the so-called “Carioca” (241 trials) and the “Black” (183 trials) (Figure 1).

As standardized by the EMBRAPA common bean breeding program, each trial is composed of 20 genotypes. The trial comprises randomized blocks with three repetitions. Tests are composed of plots of 3 m2 with 12 seeds/m. Basal fertilizers on the sowing date were applied: 240 kg/ha of monoammonium phosphate (MAP). Nitrogen (N) was applied as topdressing at V3 (third trifoliate unfold) at a dose of 45 kg/ha. Trials are divided into groups (F11, F13, and F16) that represent genotypes changes (F11 contains the same genotypes as those of 2011/2012 and 2012/2013 trials; F13 the same genotypes as those of 2013/2014 and 2015/2016 trials; and F16 the same genotypes as those of 2016/2017 trials). Finally, from each trial, we selected the following agronomic-related variables: (1) planting date (PD), days after emergence, (DAE), (2) flowering date (FD, DAE), (3) maturation date (MD, DAE), and (4) grain yield (GY, kg per ha).

As the focus of this study is to provide a broader assessment of climate variables relating them to common bean yields, considering vegetative and reproductive stages, in four (Midwest, South, Southeast, and Northeast) Brazilian production regions, we averaged genotype yields by trial. Figure S1 shows the coefficients of variation (CV) for trials across areas and seasons (supplementary information). Most CVs are below 20%, which is evidence of the representativeness of MET average values used in this study.

### 2.3 Historical farm yield

We also assessed a common bean yield official database containing municipality-level statistics (IBGE; https://sidra.ibge.gov.br/tabela/1618). The grain yield reported by IBGE, called hereafter as “farm beans yield,” was retrieved for 2011 to 2018 for all states that had cultivated common bean at least from 2011 and 2018 in different seasons (wet, dry, and winter) and regions (Midwest, South, and Southeast).

### 2.4 Environmental characterization

The agronomic variables of breeding programs were related to environmental information, such as daily climate data, by a script developed in the software R (R Core Team, 2020). The climate data set was collected from the nearest weather station to INMET (Brazilian Institute of Meteorology - https://portal.inmet.gov.br/), located at the trial state. For trials with no weather stations available in the state, we used daily climate data from NASA POWER (Sparks, 2018) and the R package EnvRtype (Costa-Neto et al., 2021b, available at https://github.com/allogamous/EnvRtype).

After relating trials to climate data, we adopted different sampling levels of environmental information (hereafter called environmental covariate – EC) to better capture the temporal variation across the crop life cycle. Each development stage was computed at a field trial level using the mean values of FD and MD observed for all genotypes evaluated in each trial. We adopted this strategy because we focused on screening EC over the common bean’s GY in four production regions in Brazil. Table 1 describes the ECs used in this study. The sampling levels adopted in this study were:

A. For crop cycle: sampling of EC in the period between planting date (PD) and harvesting, that is, involving maximum, minimum, mean, and accumulated values for each EC across the whole crop life cycle.
B. For vegetative stage: sampling of maximum, minimum, mean, and accumulated values for each EC only for the vegetative period (from PD to pre-flowering and from PD to R5 stage).
C. For the flowering period: mean sampling values for EC only for the flowering period, i.e., four days before and four days after flowering date (R6 stage), defined here as more than 50% of plants with open flowering (R6 stage).
D. For reproductive stage: sampling of maximum, minimum, mean, and accumulated values for each EC from FD to harvesting (from R6 to Harvesting)..

**Table 1.**
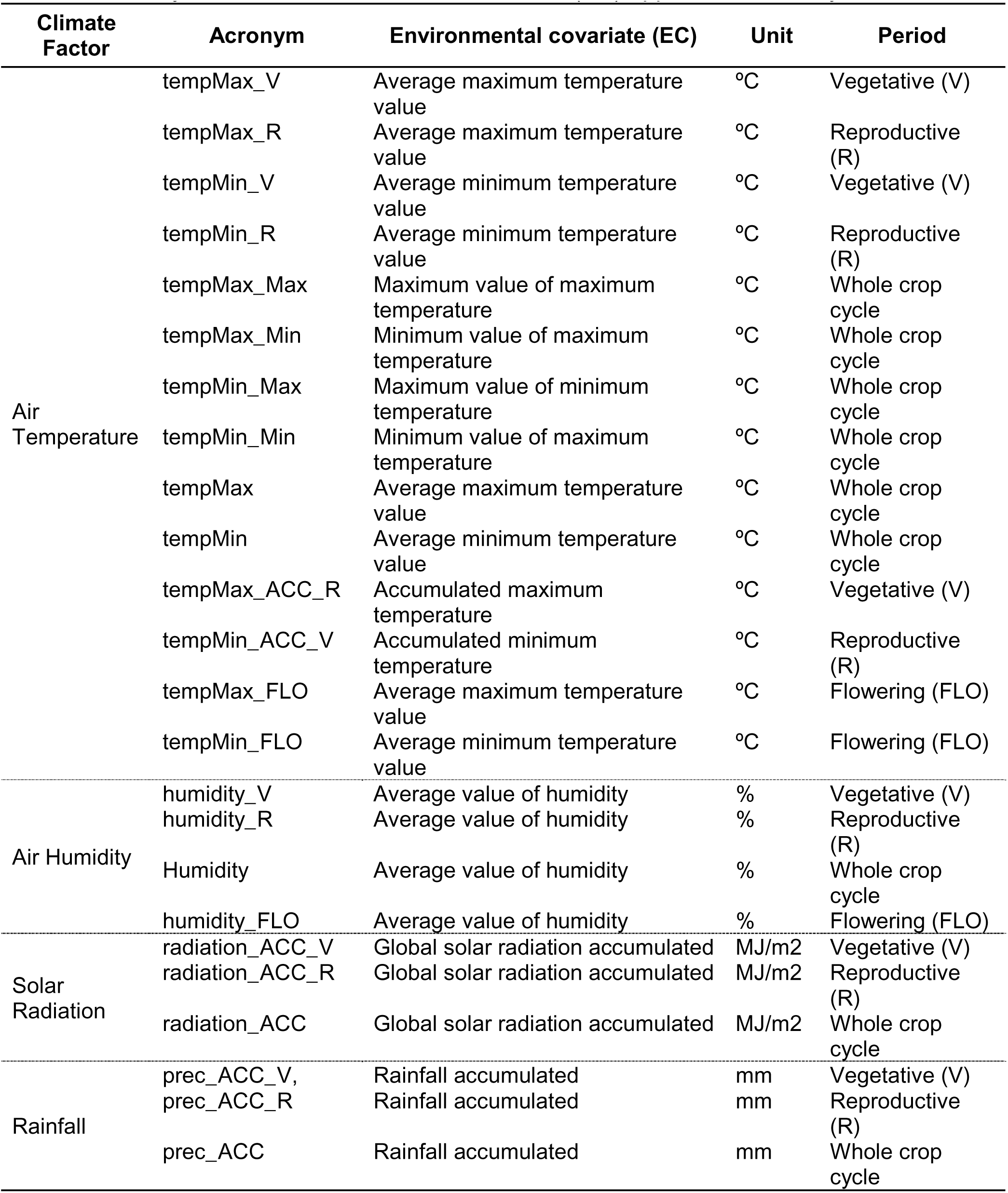
Acronyms for the environmental covariates (EC) applied in this study

### 2.5 Investigating the climate drivers of grain yield in common beans

Based on the ECs presented in the section above (Table 1), we applied and tested a Generalized Additive Model (GAM) using the mean common bean grain yield (GY) for each trial as the dependent variable. The purpose of this approach was to train the GAM models more accurately and to find the climate drivers of GY in the common bean germplasm in different production regions in Brazil. Then, we hypothesized that ECs with more ability to explain GY compose the best GAM model for each region and reveal a possible trend of climate drivers on common bean adaptation. In the next sections, we explain the mathematical description of the GAM model.

### 2.6 Generalized additive model (GAM)

#### 2.6.1 Model description

The GAM was proposed by Hastie (1986) as an alternative to the generalized linear model (GLM) (Nelder and Wedderburn, 1972), which allowed the improvement of non-parametric functions as possible predictors. In general, we represent the linear predictor of the GAM model by:

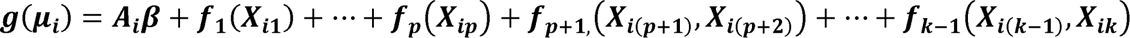

with i=1, … , n,

where 𝑔 is a specified link function; 𝐸(𝑌_𝑖_) = μ_𝑖_, with mean μ_𝑖_ and parameter vector ∅; *𝐴_𝑖_* is the matrix row of the model’s parametric components; 𝜷 is the corresponding parameter vector; and ƒ_j_ means functions with a specified parametric form or possibly specified non-parametrically or semi-parametrically (e.g., Wood, 2006).

The estimation process adopted by the GAM is analogous to the GLM (“Fisher’s Scoring”). The key difference is that the linear predictor now incorporates smooth functions of at least some (possibly all) covariates, represented as ƒ_j_, and this allows for non-linear relationships between covariates and the target variable Y. Hastie and Tibshirani (1990) described various approaches for smoothing functions, such as moving means and cubic smoothing splines. Therefore, the GAM becomes preferable than GLM when there is evidence of an unknown deterministic pattern in the data. GAM is not robust because it lacks data structure and is sensitive to non-homoscedasticity. To minimize that, we applied the GAM to each geographical region (Midwest, South, Southeast, and Northeast).

The process of selecting variables was the stepwise criterion. The fact that a covariate does not compose the selected GAM does not imply that it is not significant for the experiment. Together with pre-established covariates in the selected model, its insertion does not add significance.

#### 2.6.2 Computational implementation

All analyses in this study were conducted in the computational-statistical R environment. In particular, we used the ‘gam()’ function of the R package ‘mgcv’ (Wood, 2006). This package applies the generalized cross-validation score (GVC), which estimates the mean square prediction error based on a leave-one-out cross-validation estimation process. Finally, to select or remove covariables (in our case, ECs, as mentioned above), the stepwise method was applied based on deviance or the Akaike (AIC) criterium (Wood, 2006).

### 2.7 Predicting grain yield using research breeding database

After parametrizing the GAMs for each common bean production region (Midwest, South, Southeast, and Northeast; see Figure 1), we predicted the GY for each EC considering the trained GAM. For that, we considered as a significant EC if the p-value was lower than 5% (p ≤ 0.05). For EC variables in which the p-value was between 5 and 10%, the exclusion was based on the predictive impact of the exclusion. The predicted evaluation scenarios were performed based on the median value of numerical covariates that comprised the GAM. In this perspective, we evaluated yield in several scenarios that used the mean performance. Finally, this procedure allows us to obtain an optimal value for some climatic variables for a given season and region.

### 2.8 Predicting grain yield including farm yield historical data

Farm bean yields by municipality (item 2.3, Historical Farm Yield) were averaged for each region and year and compared with cultivars (“Black” and “Carioca”), lineages (“Black” and “Carioca”), and the highest yield lines (Lines > Quantile 0.7) of the common bean data set of the breeding program. As the farm common bean data set from IBGE is not separated by type (“Carioca” and “Black”), we aggregated the common bean types and averaged the data set by year and region. This allowed us to compare the observed farm yield (actual yield) with data sets of empirical trials of the breeding program (experimental station yield). The Northeast region was not included due to a confusion between common beans (Phaseolus vulgaris L.) and cowpea beans (Vigna unguiculata (L.) Walp) in the IBGE statistical data for the Northeast region.

## 3 RESULTS

First, we presented a nationwide overview of the distribution of grain yield (GY) and some environmental covariates (EC) in different seasons and production regions of common beans in Brazil. Then, we presented the results of the GAM in capturing the effects of EC on explaining GY. The results of this analysis are an indication of the climate-related driver GY. Next, we fitted GAMs for predictive purposes using the most significant EC found in the step mentioned above. Finally, we compared the historical farm yield with cultivars and lineages of MTE by year for a determined breeding aspect. We presented this structure for each region (South, Southwest, Midwest, and Northeast).

### 3.1 Grain yield and climate variation

Figure 2 shows the observed values of yield variation of common bean trials in different regions and seasons. There is a variation in the median (bold horizontal lines in the boxes) of common beans yield values between regions and seasons. We observed an interaction between growing regions and dry bean types. The four highest median yields were obtained in the wet season in the South region for the “Carioca” type (2,775 kg/ha ± 969 kg/ha - standard deviation) and “Black” type (2,669 kg/ha ± 1,028 kg/ha), followed by the wet season in Southeast region for the “Carioca” type (2,450 kg/ha ± 1,028 kg/ha), and the winter season in the Midwest for the “Black” type (2,417 kg/ha ± 897 kg/ha). The worst median yield was that of the Midwest in the dry season for the “Black” type (1,186 kg/ha ± 441 kg/ha) and “Carioca” type (1,275 kg/ha ± 659 kg/ha). In most cases, common beans of the “Black” type showed median yield values higher than those of “Carioca” in the wet season (Midwest, South, and Southeast). For the winter seasons, only the Midwest region had this situation. This high yield variation shown in Figure 2 is related to a climate difference (Figure 3A, B) in different seasons and regions and the level of technology applied for common bean farmers.

**Figure 2.**
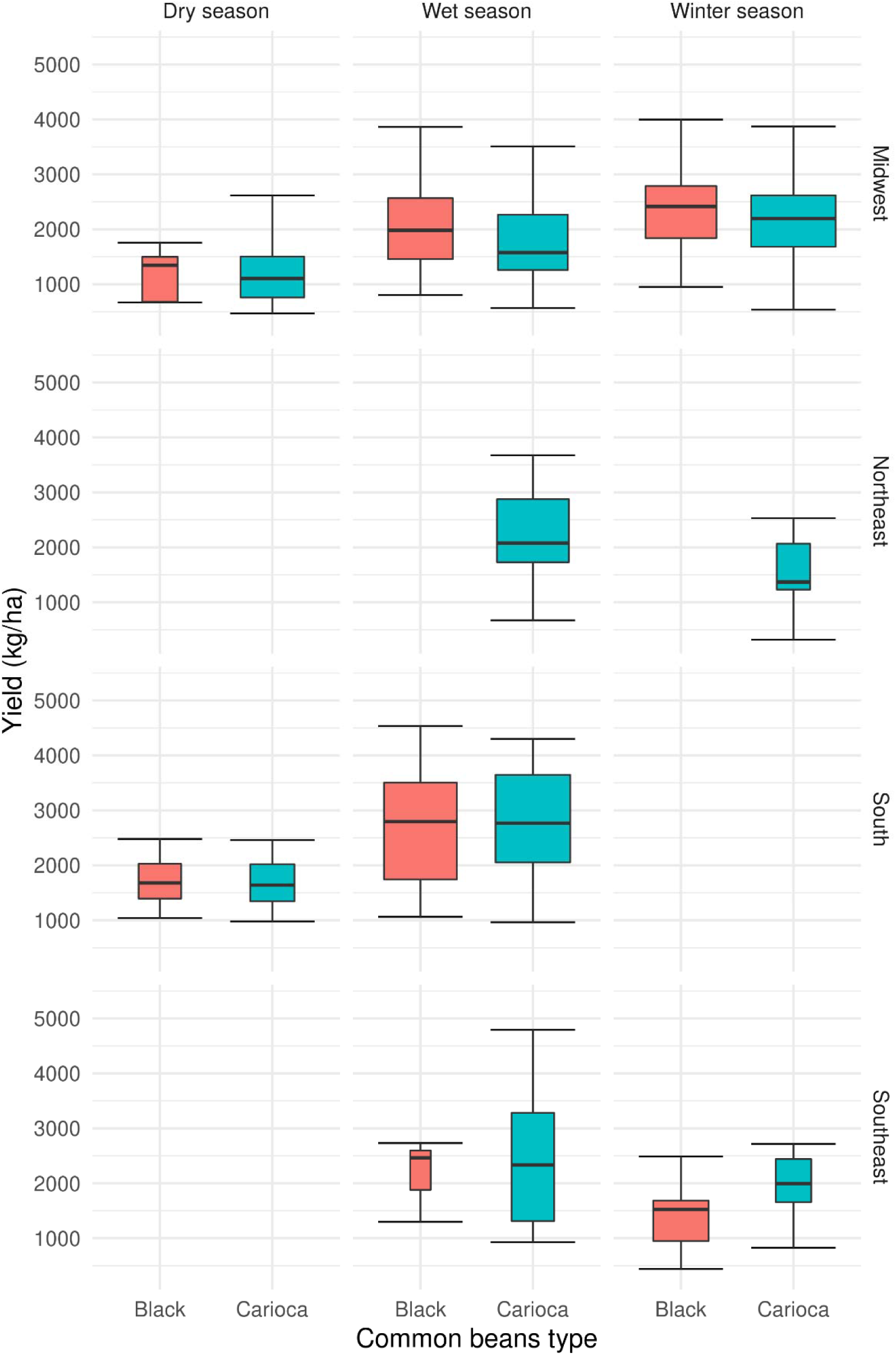
Box-plot of common bean grain yields (kg/ha) of trials in different seasons (top panel labels) and production regions (right panel labels) from 2011 to 2018. In each box-plot, the values of the boxes represent the 1^st^ quartile (25%, bottom limit), 2^nd^ quartile (50% or median, horizontal bold line) and 3^rd^ quartile (75%, top limit), and the bars indicate the minimum and maximum values observed for each distribution of yield values across different setups.

**Figure 3.**
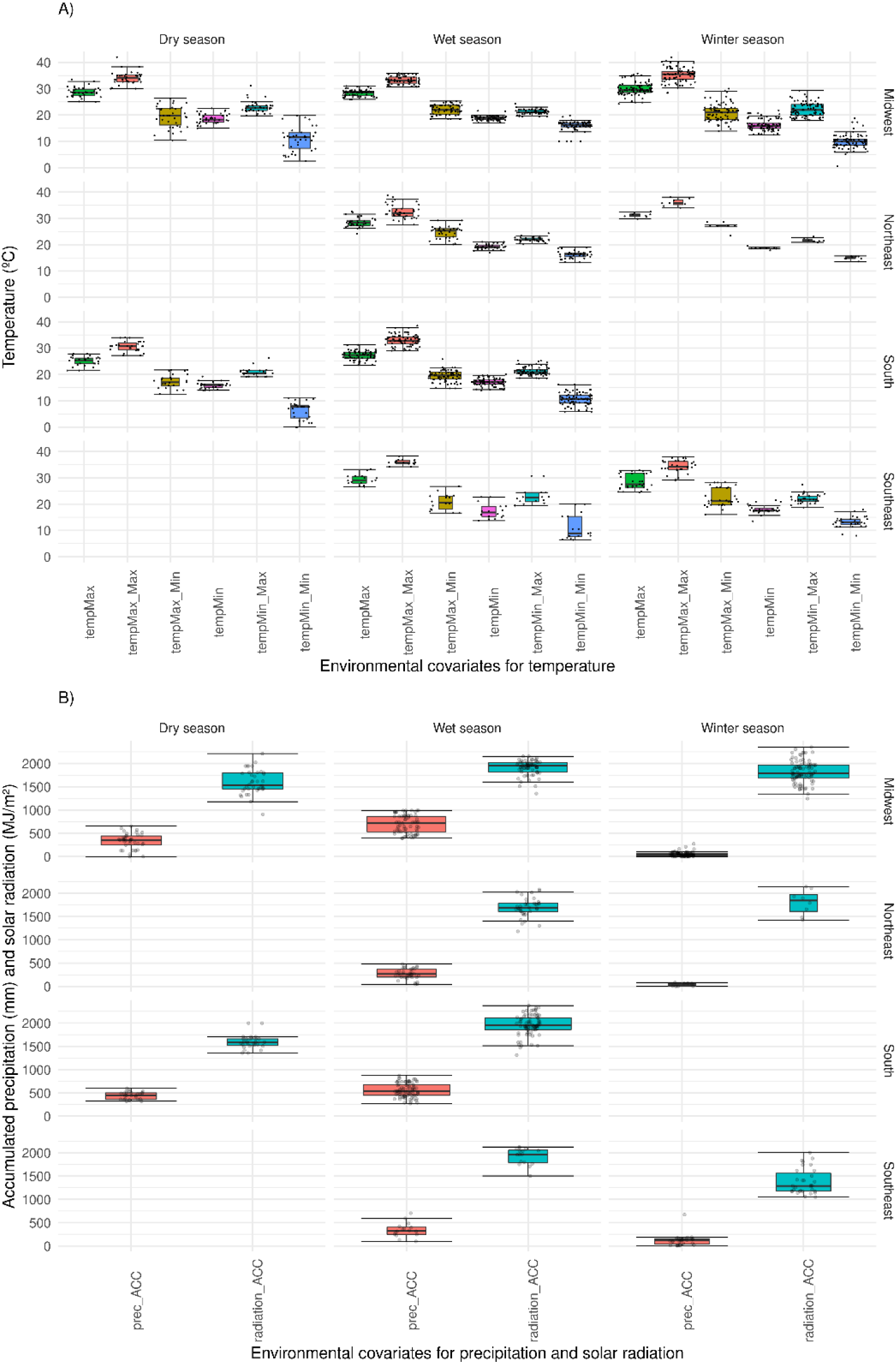
Distribution of the main environmental covariables (EC): A) temperature; B) accumulated rainfall, and accumulated solar radiation in different seasons (top panel) and production regions (right panel) for the trial experiments from 2011 to 2018. In each box-plot, the values of the boxes represent the 1^st^ quartile (25%, bottom limit), 2^nd^ quartile (50% or median, horizontal bold line), and 3^rd^ quartile (75%, top limit), and the bars indicates the minimum and maximum values observed for each distribution of climate variable values across different setups. tempMax_Max = Maximum value of maximum temperature; tempMax_Min = Minimum value of maximum temperature; tempMin_Max = Maximum value of minimum temperature; tempMin_Min = Minimum value of maximum temperature; tempMax = Average maximum temperature value; tempMin = Average minimum temperature value; prec_ACC = accumulated rainfall; and radiation_ACC = accumulated radiation.

The highest temperatures occur in the winter seasons and the lowest amplitude in wet seasons in all production regions (Figure 3A). The highest volume of rainfall occurs in the wet season for all production regions. The highest accumulated solar radiation median occurs in the wet season in the Midwest, Southeast, and South regions (Figure 3B).

### 3.2 Characterization of the South region

#### 3.2.1 Climate and categorical drivers of grain yield variation

We analyzed 101 trials in the South region. As already described in item 2.1, this region has two seasons: wet (77 trials) and dry (24 trials). These regions also cultivate two common bean types: “Carioca” (52 trials) and “Black” (49 trials). Figure S2 shows the GAM performance for this region (supplementary information). Both categorical variables, season and type, were significant at 10% and 5%, respectively. The dry season is characterized by having lower yields than the wet season does (negative estimate value in Table 2) and black type common bean showed higher yields in both seasons. This season also has the lowest values of maximum and minimum temperature, accumulated rainfall, and solar radiation (Figure 3A, B) compared to the wet season. The estimated deviance and R2 adjusted by the GAM explained 95% and 86% of the GY variation in the South region, respectively (Table 2). For the EC related to climate variables, only the maximum and minimum temperatures (tempMax_V, tempMax_R, tempMin_V, tempMin_R) and the accumulated rainfalls (prec_ACC_V, prec_ACC_R) at the vegetative and reproductive stages were significant in this region (Table 3 and Figure S3A, B, C and D, supplementary information). Among them, the maximum temperature was the most significant (p < 5%, Table 3). Maximum and minimum temperature also showed high values (24.66 and 18.44) of effective degree of freedom (edf, Table 3), which reflects the degree of non-linearity of a curve (Wood 2006). An edf equal to 1 is equivalent to a linear relationship, 1 < edf≤ 2 is considered a weak non-linear relationship, and edf > 2 means a strong non-linear relationship.

**Table 2.** Parametric coefficients of categorical variables from t test, adjusted R^2^, and explained deviance.

| Categorical Variable | Estimate | P-value | R-sq.(adj)<br>(%) | Explained<br>deviance<br>(%) |
| --- | --- | --- | --- | --- |
| **Region: South** |  |  |  |  |
| Intercept | 7.900 | < 2e-16 | 86.4 | 95.3 |
| Type "Black" | 0.068 | 0.083 |  |  |
| Dry Season | -0.885 | 0.005 |  |  |
| **Region: Southeast** |  |  |  |  |
| Intercept | 7.427 | <2e-16 | 70.1 | 84.9 |
| **Region: Midwest (all seasons: wet, dry, and winter)** |  |  |  |  |
| Intercept | 7.907 | < 2e-16 | 74.6 | 86.3 |
| Type "Black" | 0.082 | 0.098 |  |  |
| Winter Season | -0.696 | 0.008 |  |  |
| Dry Season | -0.791 | 0.000 |  |  |
| **Region: Midwest (wet and dry seasons)** |  |  |  |  |
| Intercept | 7.404 | <2e-16 | 89.4 | 95.4 |
| Type "Black" | 0.116 | 0.018 |  |  |
| Dry Season | -0.410 | 0.037 |  |  |
| **Region: Northeast** |  |  |  |  |
| (Intercept) | 7.650 | < 2e-16 | 73.2 | 86.9 |

**Table 3.**
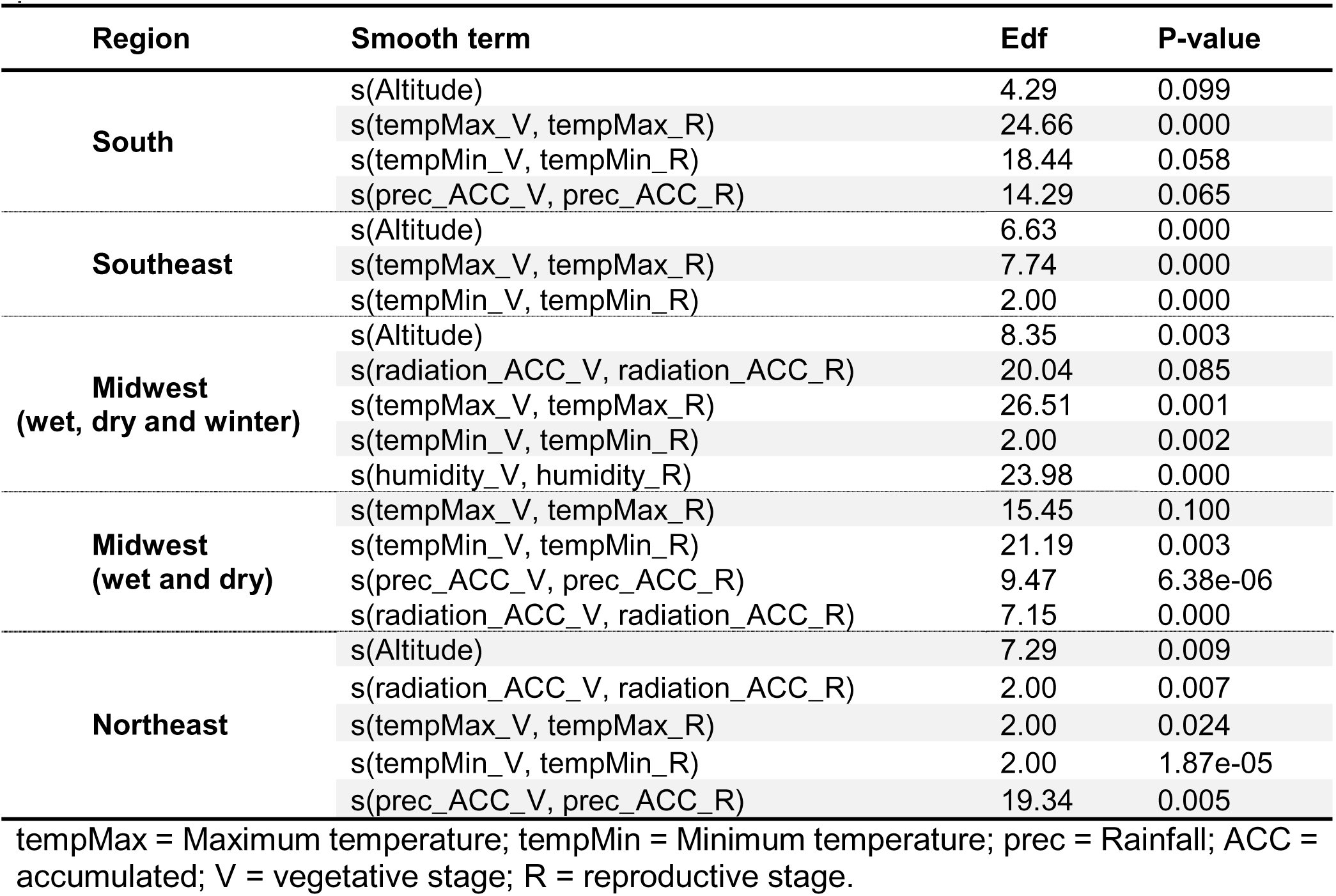
Approximate significance of smooth terms, effective degree of freedom (edf), and p-value from the F test.

In the South region, we observed maximum temperature values at reproductive and vegetative stages ranging from 20 °C to 30 °C (reproductive) and 24 °C to 30 °C (vegetative). The maximum temperature supported by the vegetative stage with no effects on yield is higher than that at the reproductive stage (Figure S3B, supplementary information). This result is in accordance with Prasad et al. (2017); on a broader developmental scale, the reproductive stage is more susceptible to heat stress than the vegetative stage. On the other hand, for the reproductive stage, only maximum temperatures higher than 26 °C exert a negative effect on yield (Figure S2B, supplementary Information). The minimum temperature ranged from 12 °C to 20 °C (vegetative stage) and 14 °C to 20 °C (reproductive stage). The reproductive stage had less sensibility to high minimum temperatures than the vegetative stage did (Figure S3, supplementary information). Minimum temperatures higher than 18 °C at the vegetative stage showed a negative effect on yield (Figure S3C, supplementary information).

Finally, the third most important climate-related EC, according to the GAM, is the accumulated rainfall at the reproductive and vegetative stages. In the South region, wet and dry seasons are rainfed systems. At both stages, accumulated rainfall ranged from 100 to 500 mm (Figure S3D, supplementary information). The reproductive stage is more sensitive to a higher volume of rainfalls. Accumulated rainfalls higher than 400 mm at this stage negatively affects yield (Figure S3D, supplementary information). Altitude was also significant at 10% (Table 3). Most trials analyzed in that region are at altitudes ranging from 600 to 800 m (Figure S3A, supplementary information).

#### 3.2.2 Environmental prediction for South region

We also predicted the GY for significant smooth terms (maximum and minimum temperature and accumulated rainfall) considering the categorical variables (season and type) (Figure 4). For that, we varied only one smooth term and fixed the other smooth terms by using the median value.

**Figure 4.**
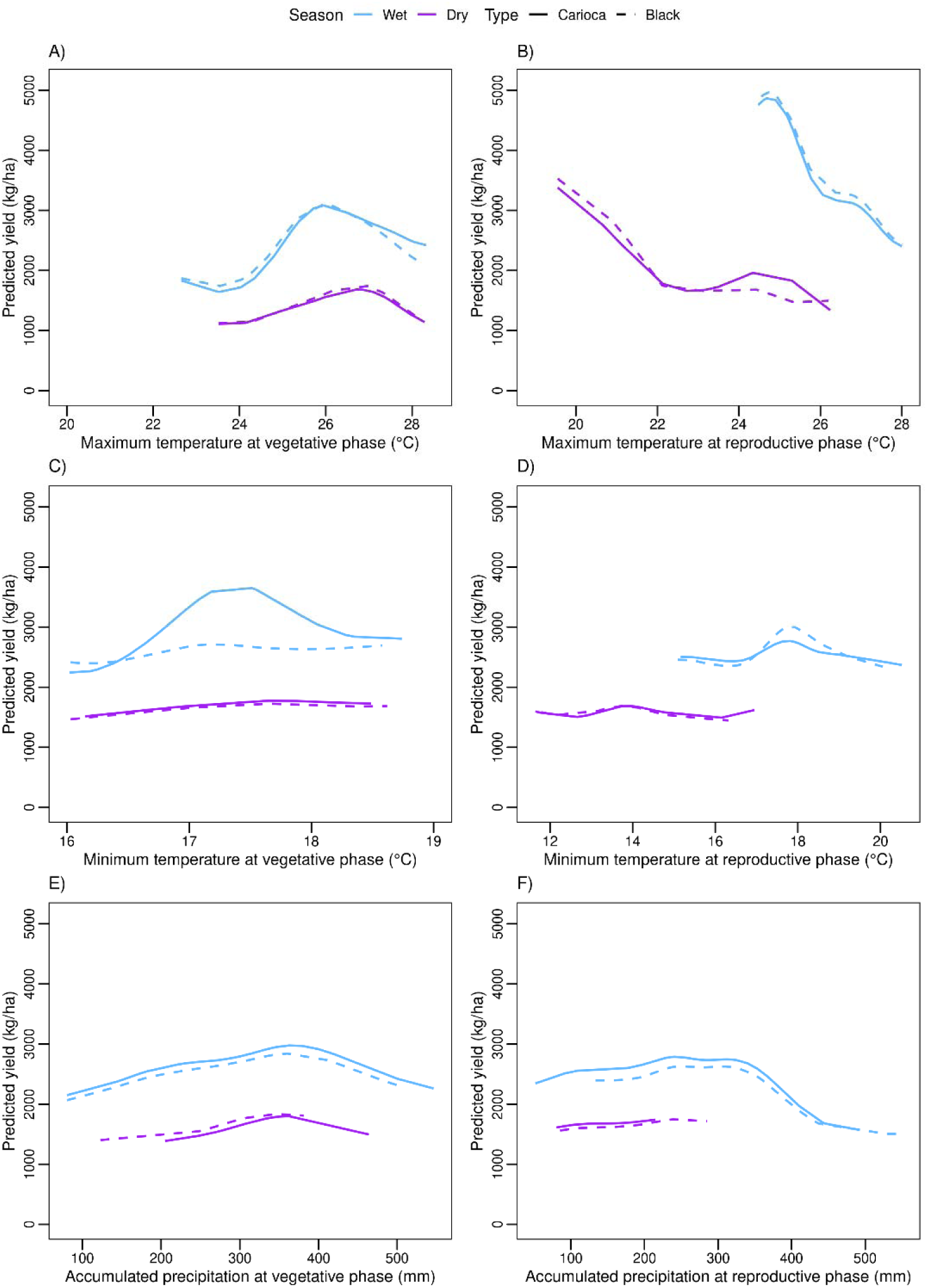
Predicted yield values for common bean growing environments in the South region as a function of the variation in the main climate drivers and key development stages: **A**) maximum temperature (°C) at the vegetative growing stage; **B**) maximum temperature (°C) at the reproductive stage, **C**) minimum temperature (°C) at vegetative stage, **D**) reproductive stage **E**) accumulated rainfall (mm) at vegetative stage; and F) reproductive stage.

The wet season showed more sensitivity than the dry season in predicting yield variations for all climate variables (maximum and minimum temperature and accumulated rainfall; Figure 4). Yield variation for both common bean types (“Carioca” and “Black”) in the wet season was higher than in the dry season (Figure 2). The common bean types “Carioca” and “Black” had similar trends for both seasons (wet and dry) and all climate variables (Figure 4A, B, D, E, and F), except for minimum temperature at the vegetative stage (Figure 4C). The “Carioca” type showed more sensitivity for minimum temperature variation at the vegetative stage than the “Black” type did. The optimum minimum temperature for the reproductive stage was 14/18 °C (wet season/dry seasons) and for the vegetative stage was 17/18 °C (Figure 4C, Table 3). Dry and wet seasons have around 50% of chance to reach the minimum optimum temperature (17/18 °C, Table 3) at the vegetative stage (Figure 5B). At the reproductive stage, wet season has around 75% of occurrence (Figure 5B) for optimum minimum temperature (18 °C, Table 3). On the other hand, the dry season reaches around 50% of the optimum minimum temperature (14 °C, Table 3).

**Figure 5.**
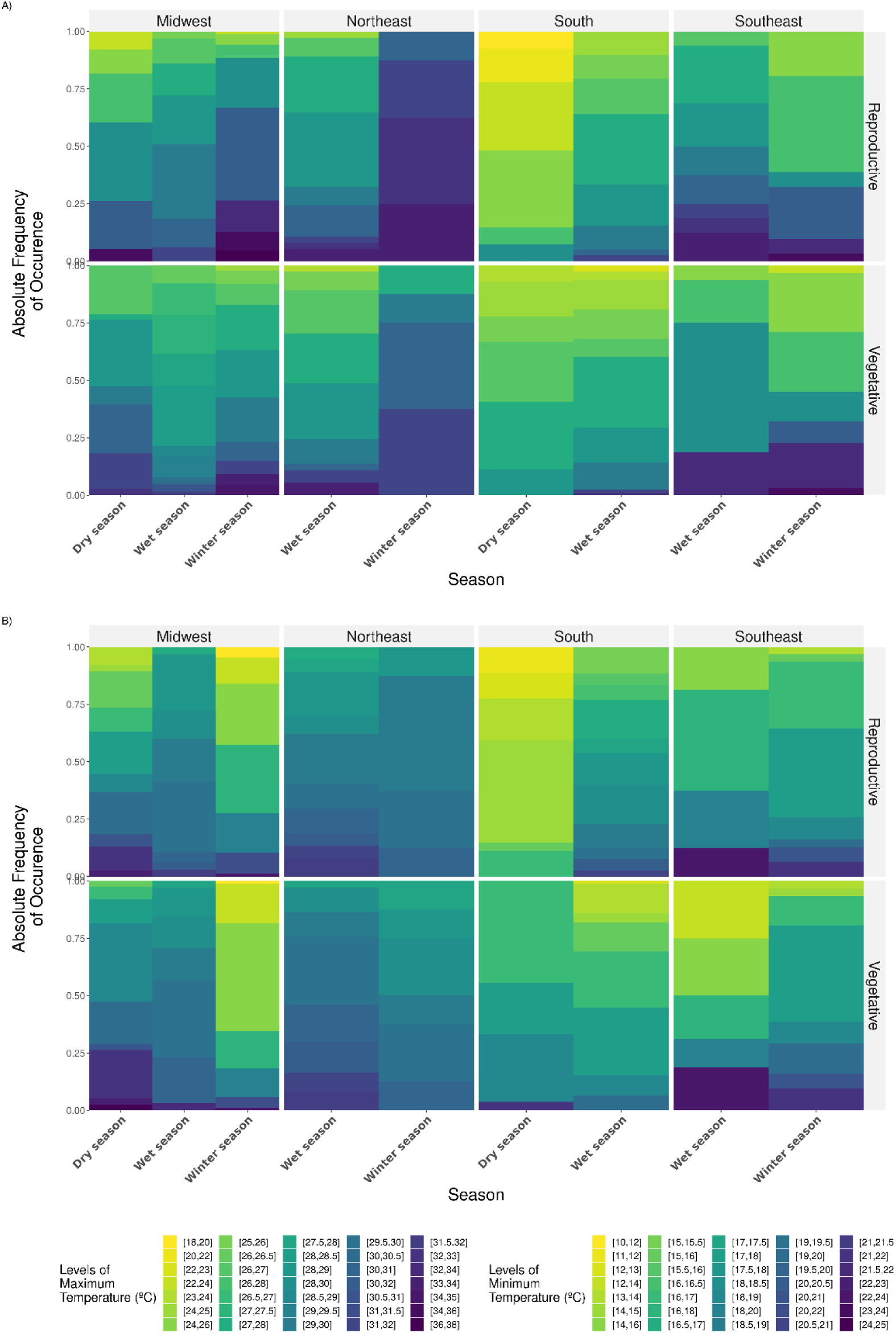
Global view of the frequency of occurrence of the main thermal-related environmental types at two development stages (vegetative and reproductive), four production regions (Midwest, Northeast, South, and Southeast) and two seasons (dry and wet). **A**) maximum air temperature and **B**) minimum air temperature.

The maximum temperature effect on predicted yield was higher at the reproductive stage (Figure 4B), corroborating the statement according to which the common bean reproductive stage is more sensitive than the vegetative stage. Optimum maximum temperature differed from wet and dry seasons at the vegetative stage (26 and 27 °C, Table 4, Figure 4A). Basically, “Carioca” and “Black” beans showed the same trend on both seasons and stages (Figure 4A and B). For the vegetative stage, both seasons (wet and dry) showed a probability of reaching the optimum maximum temperature (Figure 5A). However, for the reproductive stage, where common beans showed to be more sensitive to maximum temperatures, the wet season has higher optimum maximum temperatures than the dry season (Figure 5A). The dry season has lower maximum temperatures than the expected for the optimum maximum temperature (27 °C) for common beans in the region.

**Table 4.** Optimum values for climate variables in different regions and seasons. Max – maximum, Min – minimum, Acc – accumulated, RH – relative air humidity

| Region | Season | Climate Variable per Vegetative (V) and Reproductive (R) stage |  |  |  |  |  |  |  |  |  |
| --- | --- | --- | --- | --- | --- | --- | --- | --- | --- | --- | --- |
|  |  | Max. Temp.<br>(°C) |  | Min. Temp.<br>(°C) |  | Acc. Prec.<br>(mm) |  | Acc. Solar Radiation<br>(MJ/m<sup>2</sup>) |  | RH<br>(%) |  |
|  |  | V | R | V | R | V | R | V | R | V | R |
| South | Wet | 26 | 25 | 17-18 | 18 | 300-400 | 250-300 | - | - | - | - |
|  | Dry | 27 | 24-25 | 17-18 | 14 | 300-400 | 200 | - | - | - | - |
| Southeast |  | 30-32 | 25 | 14 | <15 | - | - | - | - | - | - |
| Midwest | Wet | 30 | 28 | - | - | - | - | >1,100 | 800 | 75 | 70 |
|  | Dry | 30 | 28 | - | - | - | - | 900-1,100 | 1,000-1,200 | 50 | 40 |
|  | Winter | 30 | 28 | - | - | - | - | >900 | >900 | 75 | 70 |
| Midwest (rainfed) | Wet | 28 | 29 | <18 | 17.5 | - | 200 | 1,200 | 1,100 | - | - |
|  | Dry | 28 | 29 | - | - | - | 200 | 1,100 | 1,000 | - | - |
| Northeast | - | - | - | - | - | 100-150 | 110 | >1,000 | >1,000 | - | - |

Optimum accumulated rainfall ranged from 300 to 400 mm (wet and dry seasons) at the vegetative stage and 250 to 350 mm (wet season) and 200 mm (dry season) at the reproductive stage (Figure 4E and F). For the wet season, higher values of accumulated rainfall showed a negative impact on yield (Figure 4F). For the vegetative stage, both seasons (wet and dry) showed a probability of having optimum accumulated rainfalls (Figure 6A). However, for the reproductive stage, the wet season has a higher occurrence of optimum accumulated rainfall than the dry season does. This season has a lower accumulated rainfall at the reproductive stage than the expected for the optimum accumulated rainfall (200 mm, Table 3, Figure 6A).

**Figure 6.**
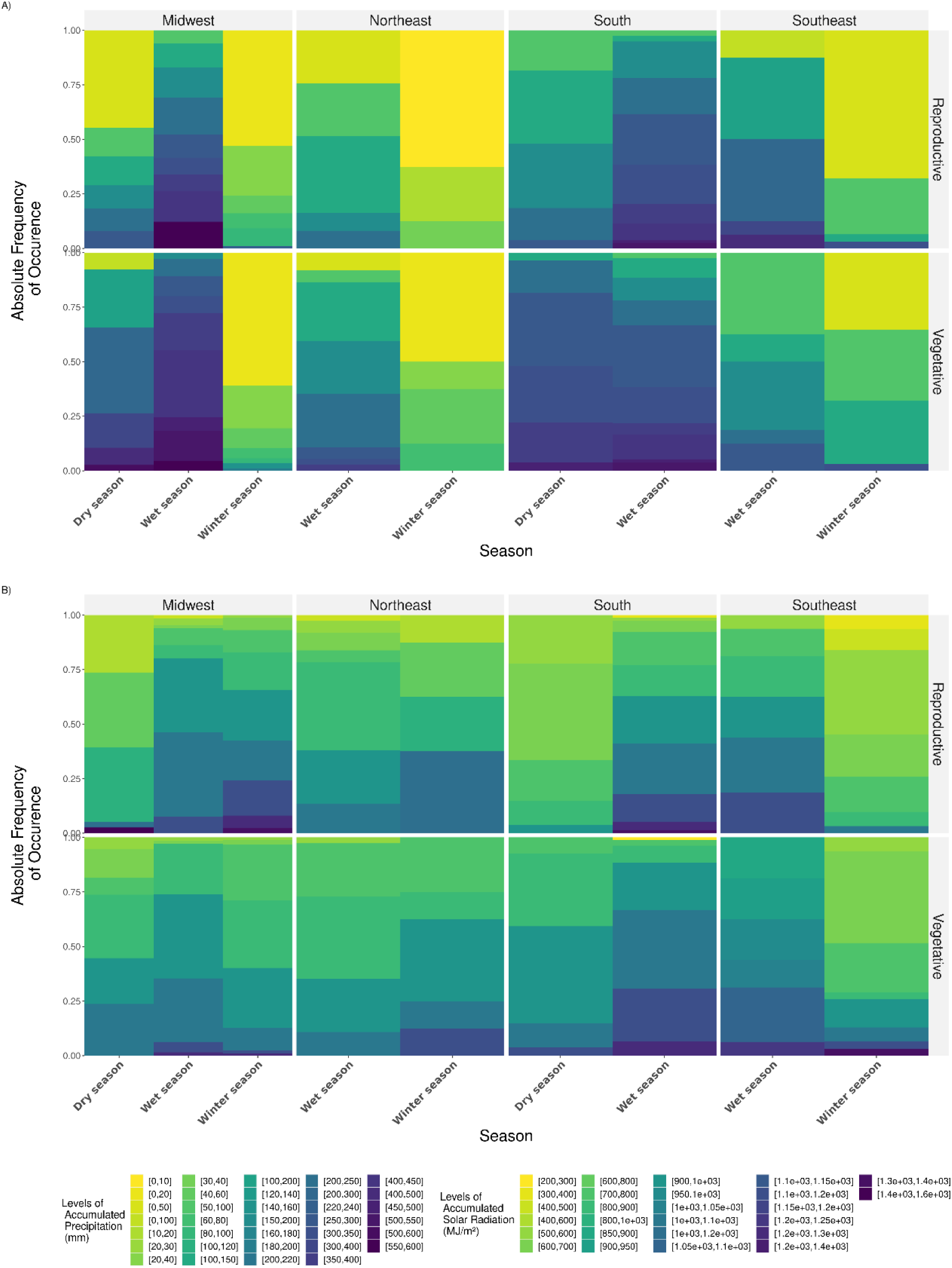
Global view of the frequency of occurrence of the main environmental types at two development stages (vegetative and reproductive), four production regions (Midwest, Northeast, South, and Southeast) and two seasons (dry and wet). **A**) accumulated rainfall and **B**) solar radiation in different regions (top panel).

### 3.3 Southeast Region

#### 3.3.1 Climate drivers affect grain yield for all types and seasons

The Southeast region comprises four States, in which we analyzed 43 trials. This region has two seasons: wet (14 trials) and winter (29 trials) (see item 2.1). They also cultivate two common beans types: “Carioca” (20 trials) and “Black” (23 trials). Figure S4 shows the GAM performance for this region (supplementary information). Differently from the South Region, here both categorical variables (season and type) were not statistically significant (p > 10%). In this region, the wet season comprises rainfed environments (no irrigation) and the winter crop is grown only in irrigated conditions.

Deviance explained by the model and adjusted R2 by GAM for the Southeast region were 85% and 70%, respectively (Table 2). Only ECs, a combination of two climate variables at two growing stages, were significant in this region: maximum and minimum temperatures (tempMax_V, tempMax_R, tempMin_V, and tempMin_R) at the vegetative and reproductive stages (Table 3 and Figure S5A, B and C; supplementary information). Maximum temperature showed the highest value (7.74) of effective degree of freedom (edf, Table 3), and maximum temperature values ranged from 24 °C to 34 °C at the vegetative stage and from 26 °C to 34 °C at the reproductive stage (Figure S5B, supplementary information). Minimum temperatures ranged from 14 °C to 22 °C at both stages, vegetative and reproductive. This region is hotter and has a higher temperature amplitude than that observed in the South region (Figure 3A). Values of minimum temperature higher than 16 °C showed a negative effect on GY at both stages (Figure S5C, supplementary information). Altitude was also significant (Table 3). Most trials analyzed in this region are located at altitudes near 0 m and from 500 m to 700 m (Figure S5A). The effects of altitude on GY are concentrated at low altitudes, from zero to 200 m (Figure S5A, supplementary information).

#### 3.3.2 Climate variable prediction for the Southeast Region

As discussed in the last subsection, there was no significant statistical difference between categorical variables (season and type) for this region. Figure 7 shows the predicted yield for smooth terms (maximum and minimum temperature). Optimum maximum and minimum temperatures for vegetative and reproductive stages were 31 °C and 25 °C, respectively (Figure 7A, B and Table 4). The vegetative stage is less sensitive than the reproductive stage for higher values of maximum temperatures (Figure 7A, B). For the vegetative stage, wet season has a higher probability of reaching the optimum maximum temperature than the winter season does (Figure 5A). However, for reproductive stage, the opposite occurred (Figure 5A). The wet season has higher yield values than the winter season (Figure 2). The increasing minimum temperature affected predicted yield at both stages, vegetative and reproductive (Figure 7C, D). However, this effect was higher at the vegetative stage. Optimum minimum temperature for vegetative and reproductive stages were 14 °C and < 15 °C, respectively (Figure 7C, D; Table 4).

**Figure 7.**
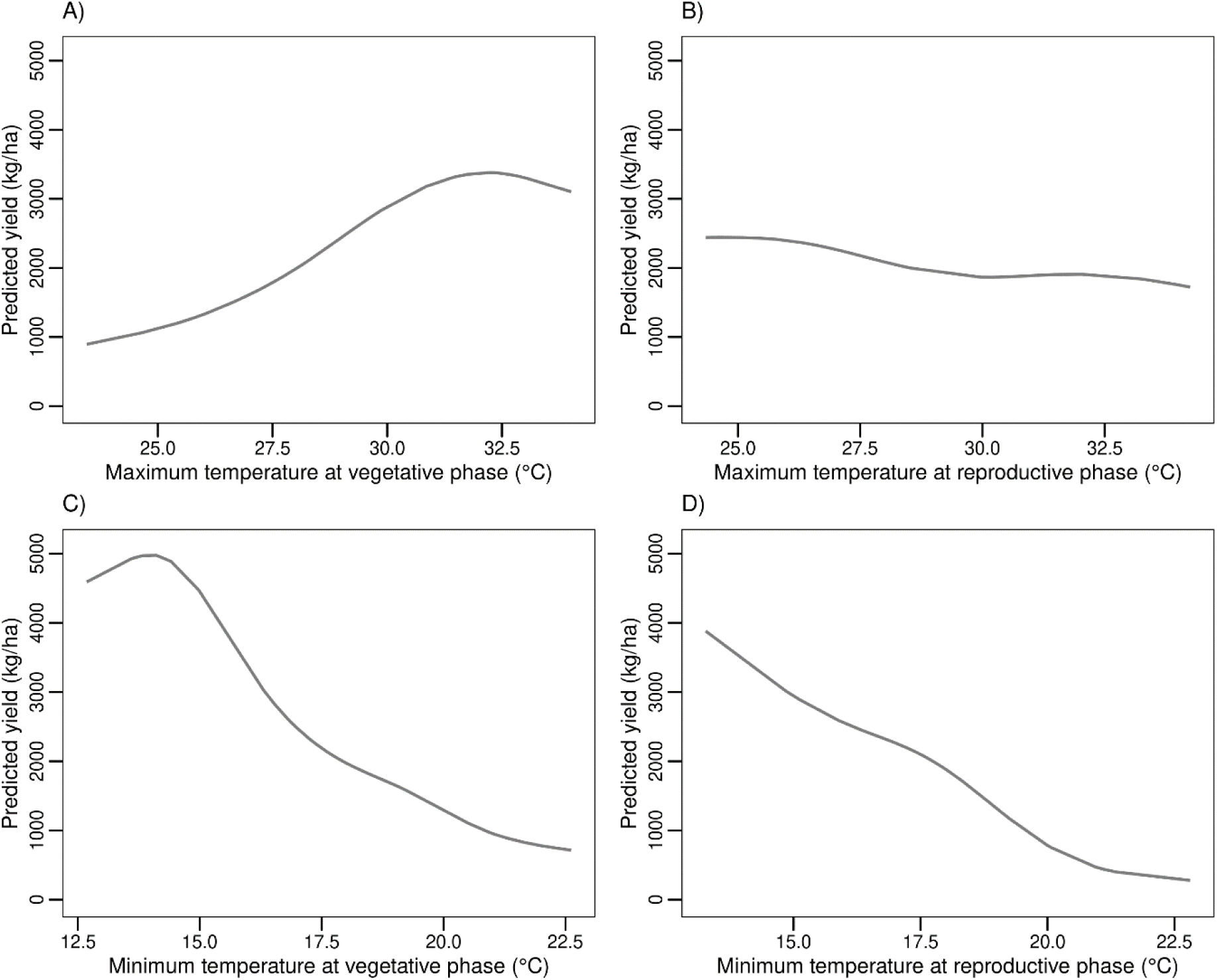
Predicted yield values for growing environments located in the Southeast region: A) maximum temperature (°C) at vegetative stage and B) reproductive stage, C) minimum temperature (°C) at vegetative stage and D) reproductive stage.

### 3.4 Midwest region

For the Midwest region, we analyzed 183 trials in four States. As we already described in item 2.1, this region has three seasons: wet (62 trials), winter (85 trials), and dry (36 trials). The state also grows two common bean types: “Carioca” (112 trials) and “Black” (71 trials). Breeding activities for common bean screening in early generation yield tests are carried out in this region at Santo Antônio de Goiás, Goiás State (latitude: −16.47 S; longitude: −49.28 W). To better understand the impacts of climate variables on common bean yield in this region, we performed two analyses considering all seasons (wet, dry, and winter seasons) and only the dry and wet seasons (rainfed seasons). This allows us to understand the importance of rainfalls for the wet and dry seasons since the winter season crop is always irrigated.

#### 3.4.1 Characterization of the Midwest region in all seasons (wet, dry, and winter)

Both categorical variables (season and bean type) were significant at 5% and 10%, respectively (Table 2). Figure S6 shows the GAM performance for this region (supplementary information). Dry and winter seasons differed from the wet season (see intercept for wet season and estimates for winter and dry seasons in Table 2). Among all seasons, the dry season showed the highest negative impact on yield (Table 2) and had the lowest common bean yields (Figure 2). The results for deviance and adjusted R2 by GAM explained 86% and 75% of the GY variation for the Midwest region, respectively (Table 2).

In this region, ECs were significant (p < 10%) in explaining GY variation in common beans. For both vegetative (V) and reproductive (R) stages, the climate variables related to air temperature (tempMax_V, tempMax_R), accumulated solar radiation (radiation_ACC_V, radiation_ACC_R), and humidity (humidity_V, humidity_R) were significant (Table 3 and Figure S7, supplementary information). The relative air humidity was the most important, followed by maximum and minimum temperatures (Table 3). Maximum temperature, humidity, and solar radiation had the highest edf values (26.51, 23.98, and 20.04, respectively; Table 3).

The maximum temperature at the vegetative and reproductive stages ranged from 25 °C to 34 °C and 24 °C to 36 °C, respectively (Figure S7C, supplementary information). The maximum temperature supported by the vegetative stage with no effects on yield was higher than at the reproductive stage (Figure S7, supplementary information). Minimum temperature showed a linear effect on GY at both stages (Figure S6C, supplementary information) and ranged from 12 °C to 22 °C. The reproductive stage was less sensitive than the vegetative stage for higher values of minimum temperature (Figure S7C, supplementary information). At the vegetative stage, minimum temperatures higher than 17 °C had a negative effect on yield (Figure S6C, supplementary information). However, the reproductive stage did not negatively affect yield at minimum temperatures in the range of the observed data set of this study (Figure 7D, supplementary information).

Solar radiation accumulated for both stages ranged from 600 MJ/m2 to 1,200 MJ/m2. In contrast, at the vegetative stage, values lower than 1,000 MJ/m2 had a negative effect on yield (Figure S7B, supplementary information). For the reproductive stage, the best range for solar radiation accumulated was 600 MJ/m2 to 800 MJ/m2. Values higher than 800 MJ/m2 negatively affect yield (Figure S7B, supplemental information).

Humidity ranged from 40 to 90% at both stages (Figure S7E, supplementary information). Moreover, at the vegetative stage, common bean cultivars may support higher humidity values than at the reproductive stage, with no effect on GY. Finally, relative air humidity values at the reproductive stage higher than 60% negatively affect the GY expression in common bean at this region. Altitude above the sea ranged from 200 m to 1000 m and its variation had a lower effect on yield (Figure S7A, supplementary information).

#### 3.4.2 Characterization of the Midwest region only in the wet and dry seasons

Considering only rainfed seasons (wet and dry), the categorical variables (season and type) were both significant at 5% (p < 5%) (Table 2). Figure S8 (supplementary information) shows the performance of the GAM for the Midwest region. Dry and wet seasons differed (see intercept for the wet season [positive] and estimate for the dry season [negative], Table 2). The GAM fitted the parameters of deviance and adjusted R2 at 89% and 95%, respectively (Table 2).

We identified eight ECs with significant effects (p < 5%) on GY variation (Table 3). The ECs were the following climate variables at both vegetative and reproductive stages: accumulated solar radiation (radiation_ACC_V, radiation_ACC_R), maximum (tempMax_V, tempMax_R) and minimum (tempMin_V, tempMin_R) temperatures, and accumulated rainfall (prec_ACC_V, prec_ACC_R) (Table 3 and Figure S8, supplemental information). Among them, rainfall was the most significant, followed by maximum and minimum temperatures (Table 3). Minimum and maximum temperatures had the highest edf values (21.19 and 15.45, respectively, Table 3).

The observed values of accumulated solar radiation ranged from 600 to 1,100 MJ/m2 and 500 to 1,100 MJ/m2 for the vegetative and reproductive stages (Figure S9A, supplementary information). For the vegetative stage, values lower than 700 MJ/m2 negatively affected the yield. However, these values are not common in this region at the vegetative stage.

Maximum temperature ranged from 27°C to 31°C (vegetative stage) and 24°C to 34°C (reproductive stage) (Figure S9B, supplementary information). Maximum temperature has a higher impact on GY at the reproductive than at the vegetative stage. For the maximum temperature range in the data set (27 to 31 °C) of this study, we did not observe impacts on yield at the vegetative stage (Figure S9B, supplementary information). On the other hand, maximum temperatures higher than 28°C at the reproductive stage had a negative impact on yield. Minimum temperatures ranged from 16°C to 22°C (vegetative) and 14°C to 22°C (reproductive) (Figure S9C, supplementary information). Minimum temperatures lower than 19 °C affected yield at the vegetative stage (Figure S9C, supplemental information).

Finally, accumulated rainfall ranged from 0 mm to 500 mm at both vegetative and reproductive stages (Figure S9D, supplemental information). However, the vegetative stage offers a higher range of rainfalls without negatively affecting the GY of common beans compared to the reproductive stage.

#### 3.4.3 Climate variable prediction for all seasons (wet, dry, and winter) in the Midwest region

For wet and winter seasons, there were no significant differences between “Black” and “Carioca” common bean types (Figure 8). Although “Black” and “Carioca” showed the same trend for all seasons, in dry season we observed that “Black” common bean was more affected by the climate variable variation than the “Carioca” type was.

**Figure 8.**
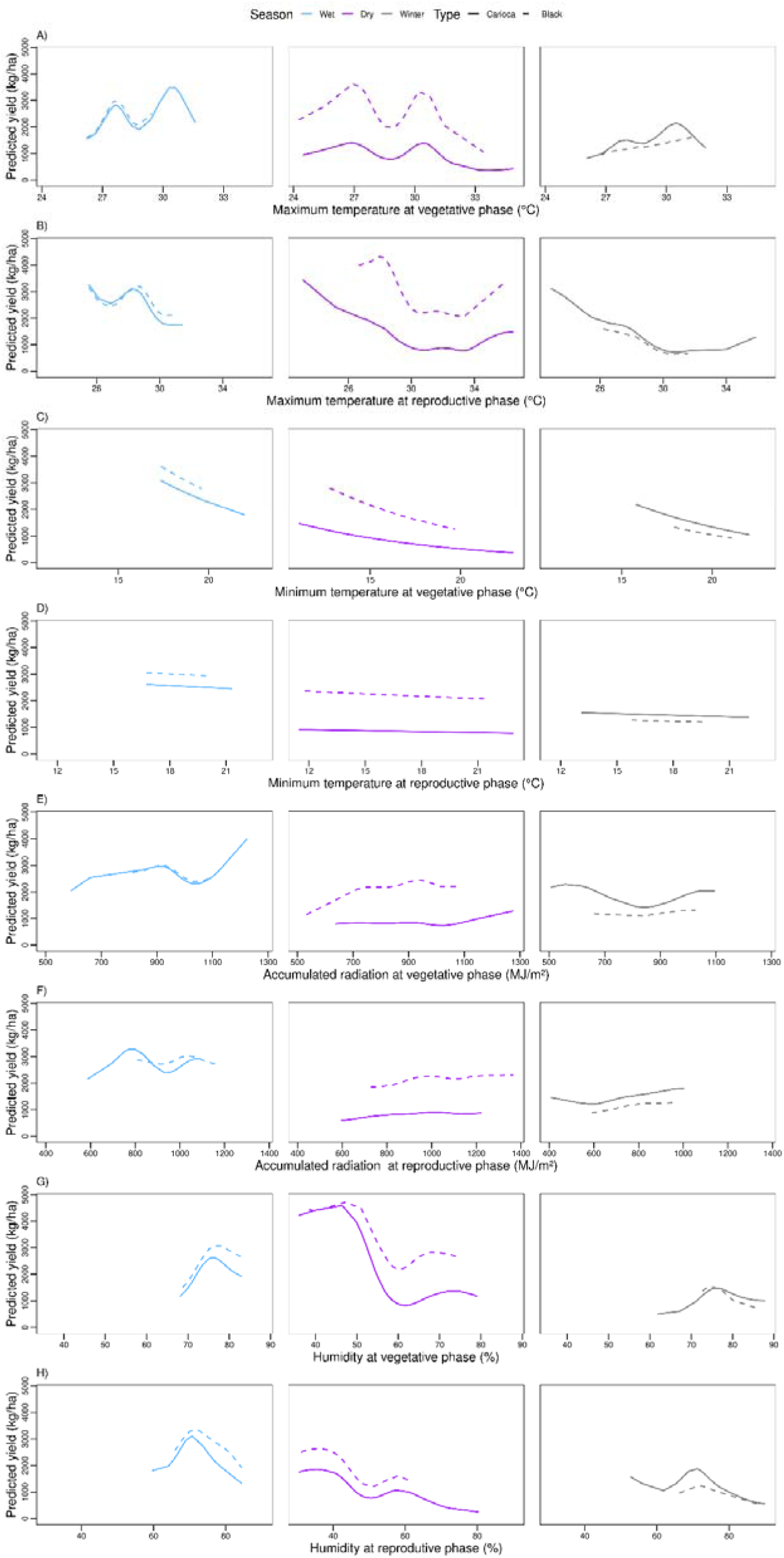
Predicted yield values for common beans in different growing environments located in the Midwest production region (wet, dry, and winter). A) maximum temperature (°C) at vegetative stage and B) reproductive stage, C) minimum temperature (°C) at vegetative stage and D) reproductive stage, E) accumulated solar radiation (MJ/m^2^) at vegetative stage and F) reproductive stage, G) humidity at vegetative stage and H) reproductive stage.

Compared to the winter season, rainfed (wet and dry) seasons have a high level of climate-driven biotic and abiotic stress and a high predominance of small-scale farms and low-input agriculture as a response to a high agroclimatic risk. On the other hand, the winter season is characterized by low biotic and abiotic (low temperatures and humidity) pressures and high levels of technology (fully irrigated and high use of fertilizer rates) (Heinemann et al., 2016).

The optimum maximum temperature for the three seasons (wet, dry, and winter) at the vegetative stage was similar as that for the Southeast region: around 30 °C (Figure 8A). The winter season has the highest probability of achieving the optimum maximum temperature at the vegetative stage (Figure 5A). Conversely, for the reproductive stage, the optimum maximum temperature was around 28 °C (Figure 8B). At the winter season, the highest probability of achieving the optimum maximum temperature occurred only at the vegetative stage (Figure 5A).

The effects of minimum air temperature on GY variation in common bean are almost linear for all seasons and both stages, vegetative and reproductive (Figure 8C,D). However, the effects of minimum temperature on the predicted yield is higher in the vegetative (higher slope angle) than in the reproductive stage. The wet season showed the most sensitive (highest slope angle) for minimum temperatures at both stages, vegetative and reproductive. In this region, minimum temperature has a low effect on GY at the reproductive stage. Finally, the highest probability of achieving an optimum minimum temperature for common beans in the winter season occurs only at the vegetative stage (Figure 5B). Thus, at the reproductive stage common beans suffer with the lowest minimum air temperatures.

Differently from air temperature, the accumulated solar radiation is the climate variable that showed the lowest significance (Table 3). Among all seasons, the highest sensibility to accumulated solar radiation occurs only in the wet season (Figure 8E, F). We observed that the optimum conditions of solar radiation for the vegetative stage was > 1,100 MJ/m2 (wet season), from 900 MJ/m2 to 1,100 MJ/m2 (dry season), and > 900 MJ/m2 (winter season) (Figure 8F). The highest probability of achieving the optimum accumulated solar radiation at the vegetative stage occurs during the wet season (Figure 6B). Conversely, for the reproductive stage, the highest probability of achieving the optimum solar radiation conditions at the reproductive stage occurs in the winter season, followed by the wet season (Figure 6B). The optimum values for accumulated solar radiation were ∼800 MJ/m2 (wet season), from 1,000 MJ/m2 to 1,200 MJ/m2 (dry season), and > 900 MJ/m2 (winter season) (Figure 8F).

Air humidity (%) is the most significant climate variable for the Midwest region in all seasons (Table 2). Common bean germplasm is more sensible to higher humidity values at the reproductive than at the vegetative stage (Figure 8G,H). For the vegetative stage, humidity showed a higher effect on GY in the dry season, followed by the wet season (Figure 8G), both conducted under rainfed growing conditions. We observed that the optimum humidity for the vegetative stage was 75% for both seasons, wet and winter. However, for the dry season, this optimum value was 50% (Figure 8G). Finally, for the reproductive stage, the optimum humidity at the reproductive stage ranged from 40% (dry season) to 70% (wet and winter seasons) (Figure 8H).

#### 3.4.4 Climate variable prediction for wet and dry seasons in Midwest region

As observed in the subsection (3.4.2), there was difference between “Black” and “Carioca” common bean types and seasons (wet and dry). The wet season showed more sensitivity for climate variation than the dry season (Figure 9), except for maximum air temperature (Figure 9B,C).

**Figure 9.**
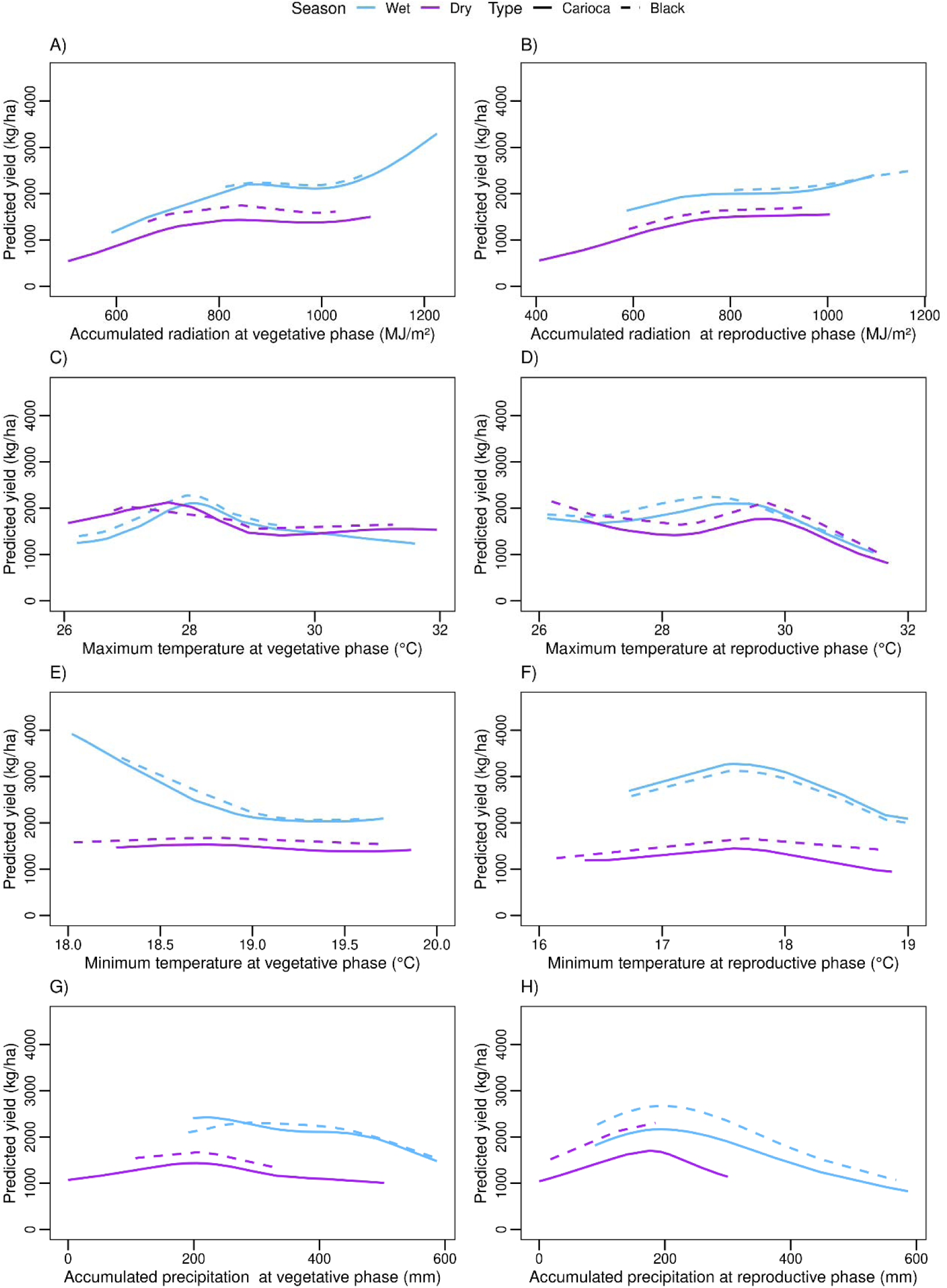
Predicted common bean yield for the Midwest region in wet and dry seasons: A) accumulated solar radiation (MJ/m^2^) at vegetative stage and B) reproductive stage; C) maximum temperature (°C) at vegetative stage and D) reproductive stage, E) minimum temperature (°C) at vegetative stage and F) reproductive stage, G) accumulated rainfall (MJ/m^2^) at vegetative stage and H) reproductive stage.

As expected due to the common bean ecophysiology, our results point that the vegetative stage demands more radiation than the reproductive stage (Figure 9A,B). The optimum values for accumulated radiation achieved were 1,200 MJ/m2 (wet season) and 1,100 MJ/m2 (dry season) for the vegetative stage; and 1,100 MJ/m2 (wet season) and 1,000 MJ/m2 (dry season) for the reproductive stage (Figure 9A,B). Among these seasons, the higher probability to reach the optimum accumulated radiation is at the wet season, which was expected, concentrated during the summer season (Figure 5B).

During the wet and dry seasons, the maximum temperature ranged from 26 °C to 32 °C at both the vegetative and reproductive stages (Figure 9C). Basically, for the vegetative stage, the variation in maximum temperature in Midwest region has no effect on GY in both seasons. On the other hand, for the reproductive stage, only values of maximum temperature higher than 30 °C have a negative impact on common bean production (Figure 9D). The optimum maximum temperature for the vegetative stage is around 28 °C in both seasons (Figure 9C), and for reproductive stage this value is 29 °C in both seasons (Figure 9D). During the vegetative stage, the dry season has a higher probability of reaching optimum conditions of maximum temperature. However, for the reproductive stage, the wet season has a higher probability of reaching these optimum conditions for common bean production (Figure 5A,B).

The minimum temperature ranged from 16 °C to 20 °C and did not affect yield in the dry season at both stages, vegetative and reproductive (Figure 9E,F). On the other hand, in the wet season, minimum temperatures higher than 18 °C affect the yield negatively at the vegetative stage (Figure 9E). At the reproductive stage, the optimum minimum temperature is 17.5 °C (Figure 9F). The wet season has a high probability of reaching the optimum minimum temperature at both stages (Figure 5B).

The wet season showed more sensibility to accumulated rainfall (Figure 9G,H). The accumulated rainfall has a higher impact at the reproductive stage than at the vegetative stage (Figure 9G,H). The optimum accumulated rainfall at the reproductive stage was 200 mm in both seasons (Figure 9G,H). For the vegetative stage, both seasons has a high probability of reaching the optimum accumulated rainfall (Figure 6A). However, at the reproductive stage, the wet season has a higher probability of supplying the accumulated rainfall demand for this stage (Figure 6A).

### 3.5 Northeast region

#### 3.5.1 Characterization of the Northeast region

The Northeast is the last region under analysis in this study. We used information from 37 field trials in the wet season (rainfed conditions) and seven trials in the winter season (irrigated conditions). In these 44 trials analyzed, only “Carioca” type was cultivated. Fig. S10 shows the GAM performance for this region (supplementary information). Our results show that categorical variables (season and type) were not statistically significant for this region (p > 10%) (Table 2).

The explained deviance and the adjusted R2 by GAM at the Northeast region was 87% and 73%, respectively (Table 2). We identified nine ECs at two key development stages, as already observed in other regions (vegetative and reproductive). They involve temperature-related factors (tempMin_V, tempMin_R, tempMax_V and tempMax_R), solar radiation (radiation_ACC_V, radiation_ACC_R), and rainfall (prec_ACC_V, prec_ACC_R) (Table 3 and Figure S11 A,B,C,D,E, supplementary information). The most significant climate variable was minimum air temperature (tempMin_V and tempMin_R), followed by the accumulated rainfall (prec_ACC_V and prec_ACC_R) (Table 3). Rainfall had the highest edf value (19.34, Table 3).

Accumulated radiation and maximum and minimum temperatures showed a linear effect on GY (Figure 10B,C,D supplemental information). For these ECs, minimum temperature in the Northeast region had the same range at the vegetative and the reproductive stage. At the vegetative stage, the minimum air temperature ranged from 18 °C to 21 °C, almost equal to the reproductive stage, which ranged from 17 °C to 21 °C (Figure S11D, supplementary information). Minimum temperatures lower than 21 °C affected the yield at the vegetative stage. For the reproductive stage, the same situation happens for minimum temperatures lower than 20 °C. Maximum temperatures at both stages ranged from 24 to 32 °C (Figure S11C, supplementary information). For the vegetative stage, this range of maximum temperatures does not affect yield. In the other hand, for the reproductive stage, maximum temperatures higher than 29 °C have a negative effect on yield (Figure S11C, supplementary information).

**Figure 10.**
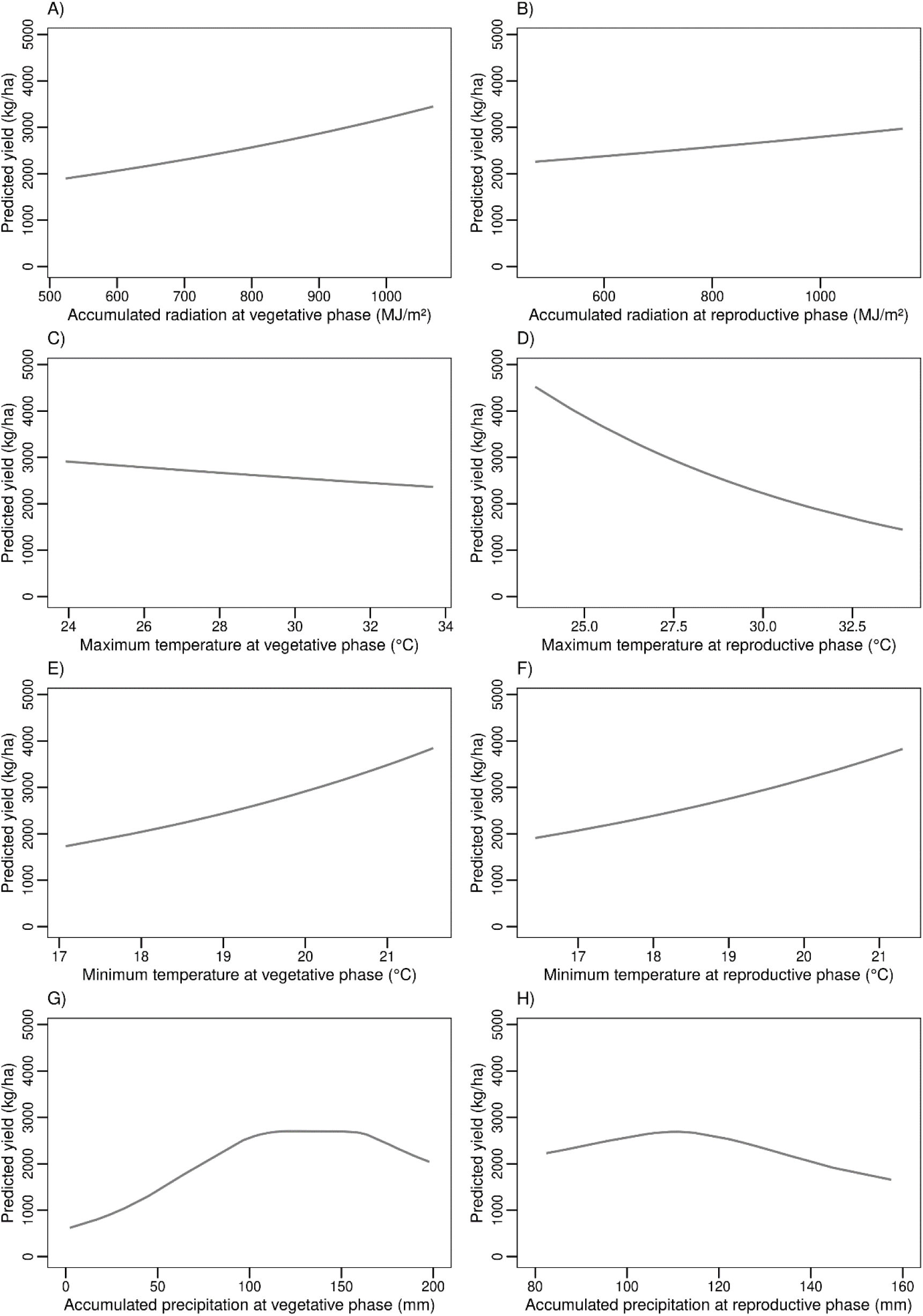
Predict common bean yield for Northeast region: A) accumulated solar radiation (MJ/m^2^) at vegetative and B) reproductive stage; C) maximum temperature (°C) at vegetative and D) reproductive stage, E) minimum temperature (°C) at vegetative, F) reproductive stage, G) accumulated rainfall (mm) at vegetative and H) reproductive stage.

The accumulated rainfall (prec_ACC) values ranged from 25 mm to 350 mm at the vegetative stage, and from 50 mm to 250 mm at the reproductive stage (Figure S11E, supplementary information). At both stages, vegetative and reproductive, accumulated rainfall lower than 100 mm has a negative impact on yield. Solar radiation (radiation_ACC) has also an important effect on common bean yield in the Northeast region. The observed values for accumulated radiation ranged from 600 to 1,000 MJ/m2 and 500 to 1,100 MJ/m2 for the vegetative and reproductive stages, respectively (Figure S11B, supplementary information). For both stages, accumulated radiation lower than 1,000 MJ/m2 has a negative effect on yield. Finally, we observed a positive effect of altitude on GY in field trials located between 300 m to 400 m above the sea level (Figure S11A, supplementary information).

#### 3.5.2 Climate prediction for the Northeast region

In the Northeast region, categorical variables, season, and bean types (“Carioca”, “Black”) were not significant (p > 10%) (Table 2). Figure 10 shows the predicted GY conceived for smooth terms due to the main ECs described in the last subsection: maximum and minimum temperature, accumulated rainfall, and accumulated radiation. A positive trend of GY occurs due to increased values of accumulated radiation at both vegetative and reproductive stages (Figure 10A,B) although the impacts of this climate factor were higher at the vegetative than the reproductive stage (higher slope in Figure 10A than in 10B). As common bean production correlates with the increase in solar radiation, we observed that the optimum radiation to achieve potential yield values occurs in environments with at least 1,000 MJ/m2 accumulated at the vegetative and reproductive stages (a total of ∼2,000 MJ/m2 at both stages).

Maximum temperature showed a negative trend for both stages. The higher the maximum temperature, the lower the GY (Figure 10C,D). The impact of maximum temperature was higher on GY at the reproductive stage (Figure 10D, higher slope). Due to the linear trend of predicted GY at both stages, it is not possible to determine the optimum climate limits.

Minimum temperature has a linear positive trend for both stages. The higher the minimum temperature, the higher the GY (Figure 10E,F). The impacts of minimum temperature were quite similar at both stages. Due to the linear trend on predicted GY in both stages, it is not possible to determine the optimum climate limits. However, combining our maximum and minimum temperature results, we infer that the optimum thermal conditions for common bean adaptation and reaching the potential yield are not frequent in this environment mainly in the winter season.

Finally, our results suggest that the optimum limit for accumulated rainfall ranges from 100 to 150 mm and 110 mm at the vegetative and reproductive stages, respectively (Figure 10G,H).

### 3.6 Breeding perspectives for the Midwest, South, and Southwest regions

A different way of analyzing adaptation results of a breeding program for a specific region is by comparing farmer yield (actual yield, described in item 2.8 above; Figure 11, blue line), cultivars (“Black” and “Carioca” types; Figure 11, green line), lineages (“Black” and “Carioca” types; Figure 11, orange line), and the highest yield lines (Lines > Quantile 0.7; Figure 11, black line) in different regions and seasons by year. The closer the farmer yield is to cultivars, lineages, and the highest yield lines, the more difficult it would be to increase farm yield without explicitly addressing cultivar adaptations to a specific region. Farmer yield, also called “actual yield,” is generally below cultivar, lineage, and highest yield lines. In different regions and seasons, farm yield increases or remains relatively constant through time, reflecting better management practices (Figure 11). On the other hand, cultivar, lineage, and high yield lines tend to decrease in most regions and seasons, except for the wet season (Midwest region) and the winter season (Southeast region), likely reflecting a lack of adaptation to season/region (Figure 11). For most seasons/regions, farm yield became closer to cultivar, lineage, and high yield lines: Midwest (dry and winter season), South (dry season), and Southeast (winter). Notably, for the Midwest, the winter season, characterized by having farmers with a high level of technology (i.e., infrastructure suitable for mechanical cultivation, irrigation by center pivot, application of high levels of nitrogen rates, receptiveness to new technologies, and common bean cultivation at commercial scales), the farmer yield is closer (inside a confidence interval) to cultivar, lineage, and high yield lines. Conversely, for wet seasons and all regions, farmer yield is the furthest away from reaching cultivar, lineage, and high yield lines. Unlike the Midwest region/winter season, an adapted cultivar is not limiting to an increased yield, and investments in better management practices may lead to increased yields.

**Figure 11.**
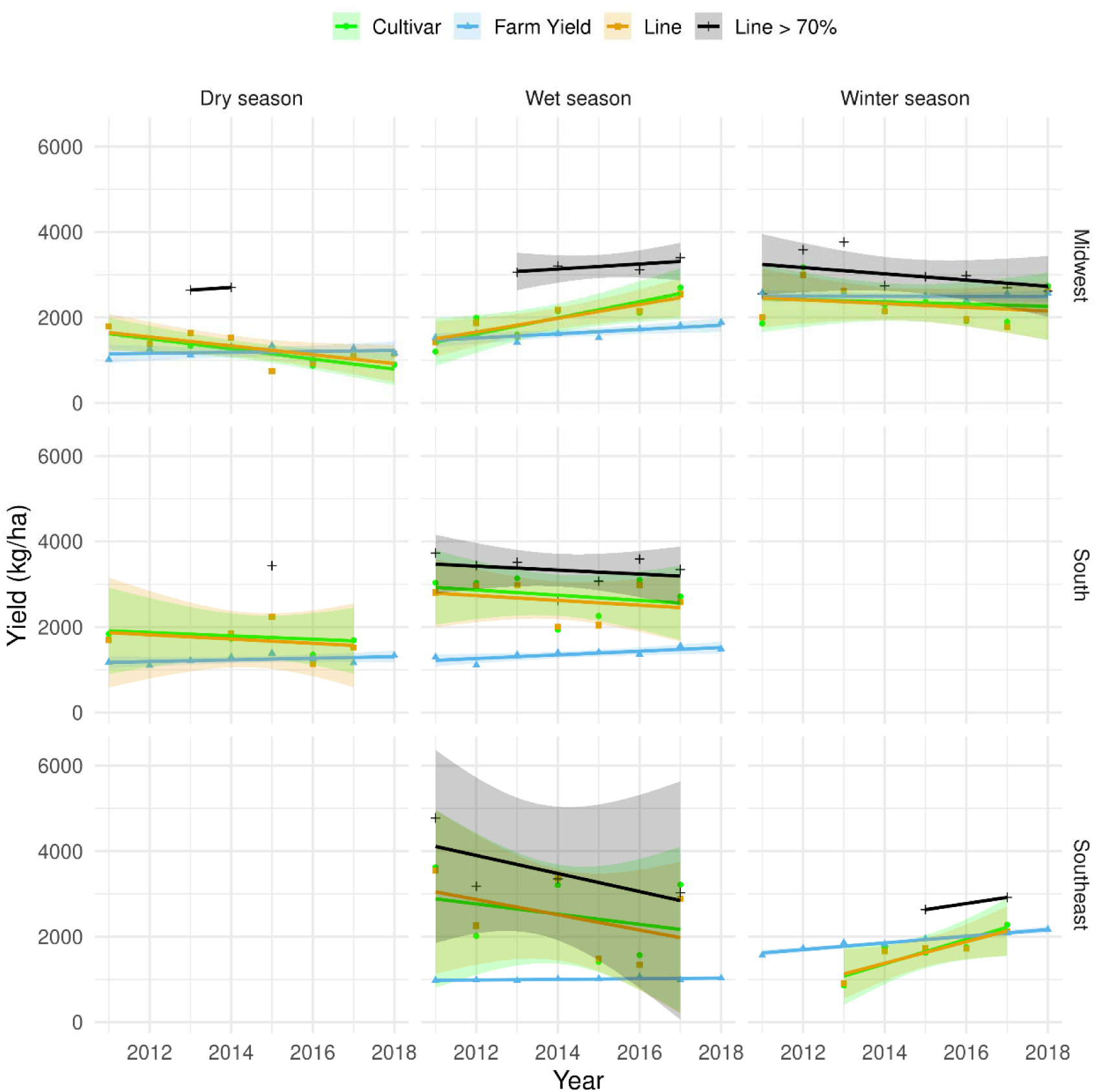
Linear regression for farm yield (blue line), cultivar (green line), lineages (orange line), and high yield lineages (Lineages > Quantile 0.7; black line) in different seasons (top panel) and regions (right panel) over the years. Lines represent the linear regression and bands represent the confidence interval.

## 4 DISCUSSION

In this study, we presented the first report on (1) the modeling of the association between climate and phenotypic variation based on design prediction models for large growing regions, (2) an effective approach to characterize growing environments using GAM as an exploratory and predictive tool; and (3) an application of the so-called “enviromics prediction” (Resende et al., 2021; Costa-Neto et al., 2021b) for GY in common beans and its consequences in developing climate-smart varieties. In the context of enviromics prediction, here we discuss first the use of GAM in the hereafter named “climate prediction” for determining possible drivers of yield variation across extensive and complex target production regions, such as in common beans growing regions in Brazil. Then, we discuss the implications of the main climate drivers found in this study in the current breeding program pipeline. Finally, we envisage future opportunities and challenges faced by research and breeding frameworks.

### 4.1 Usefulness of GAM in defining climate limits of adaptation and breeding zones

The model selection process used in this study ensured that goodness-of-fit was balanced in face of model complexity by adding explanatory variables one at a time only if they added to the model’s predictive ability. This process showed that, for categorical variables, season (water, dry, and winter) and type (“Carioca” and “Black”) are important only for the South and Midwest regions. The climate variable maximum and minimum temperatures are important for all regions. Accumulated rainfall is significant only for the South, Midwest (rainfed system: wet and dry seasons), and Northeast regions, while accumulated solar radiation is significant only for the Midwest and Northeast region in low latitude locations (7° S - 16° S). Based on the predicted yields by GAM (described in item 2.3.2), we obtained the “optimum” values of climate variable for different seasons and regions (Table 3). We observed that “optimum” climate variables varied across seasons and regions. Then, we could infer the limits of adaptation for common beans in each combination of region, season, and sometimes grain type. In terms of breeding, the limits highlight the importance of capitalizing the genetic x environment (GE) interactions in the selection process; in other words, the breeding process should be within and specific to each mega-environment or market concept and not to a wide adaptation.

### 4.2 From the farm to the plant breeding research

We also predicted the GY at farm level (considering a core of genotypes at a given environment). In the next sections, we apply the conclusions from the farm to breeding. The potential yield of a certain region depends on environmental factors (E) and genetic factors (G) related to the core of genes capable of good responses to E variations. The breeding program creates diversity of G by selecting adapted genes, testing them in a diverse set of growing conditions, and finally launching a new cultivar.

Based on our approach, we could visualize the use of climate prediction focused on understanding E for a pool of genotypes. This can also be used for particular genotypes to study reaction-norm (linear responsiveness of genotypes to a certain environmental factor) and gene-based cause of adaptability and yield stability in a diverse set of growing conditions (which makes the selection and targeting of novel G difficult). The differences in gene-based reaction norms for different genotypes led to the phenomena called “genotype-by-environment interaction” (G×E). A common use of environmental factors to understand E and G×E relies on factorial regression (e.g., Wood, 1976; Denis, 1980; Costa-Neto et al., 2020) and partial least squares (e.g., Voltas e al., 1999; Monteverde et al., 2019; Porker et al., 2020). As Arnold et al. (2019) suggested, the common use of reaction-norm as a linear response to a certain environment gradient may not be the best alternative to shape the actual phenotypic plasticity observed in field crops. Other approaches using non-linear methods, mostly considering polynomial responses or non-parametric responses (e.g., GAM), might describe better the reaction-norm responses and consequently find covariates affecting the phenotypic plasticity of a certain trait (Arnold et al., 2019).

Here, we envisage that GAM-based climate prediction can be a powerful tool to support the design of crop ideotypes for each region, season, and future scenario, leading breeding programs to develop more adapted varieties. For instance, as observed for some regions such as the Northeast, the effects of high temperatures (resulting in heat stress) may hinder future genetic gains and common beans yields. Therefore, breeders can use this information to screen the reaction of candidate cultivars in early trials conducted under multi-environment conditions to achieve suitable favorable responses, for instance the observed thermal conditions of some particular region (or season).

### 4.3 Usefulness of climate prediction to fill breeding gaps in research on common beans

In plant breeding, large-scale environmental data is useful in explaining the environmental drivers of variations in key traits (e.g., grain yield). Here, we provided the first report of the use of GAM with such a purpose and then discussed how GAM-based climate prediction may fill knowledge gaps in breeding decisions for common beans.

The major common beans breeding program in Brazil, led by EMBRAPA, has a wide adaptation in all regions (South, Southeast, Northeast, and Midwest), with a focus mainly on GY improvement. Consequently, other important traits are selected due to indirect factors related to pleiotropy and epistasis and during the cultivar testing stage, where it is possible to verify the resilience of elite cultivars to a core of complex diseases and managements (e.g., irrigation). However, a limitation of this strategy is the fact that it does not consider limitations in different regions and seasons. Also, the common beans breeding program does not consider climate variables in its analysis.

There is a gap in the understanding of the actual importance of environmental factors to phenotypic variation and G×E interaction for GY (see Table S1, supplementary information). Pereira et al. (2017) present an extensive analysis involving 16 elite cultivars of the “Carioca” type tested across 62 environments in over 30 locations across the Midwest and Southeast regions of Brazil. Despite the discussions made in that study, the authors analyzed only variance components related to genetic variation (G), environmental variation (E), and G×E.

The phenotypic variation in GY is ∼2% partly due to G causes, from 82% (Midwest) to 88% (Southeast) due to E-related causes, and from 10% (Midwest) to 16% (Southeast) due G×E. Rocha et al. (2020) evaluated 15 elite cultivars of the types “Carioca” and “Black” over four years in Northeast Brazil. Although these authors used GGE analysis (focused on G and G×E), we recall again the variance component table of this study to verify the importance of G, E, and G×E. A major importance of E-related factors was also observed: for the “Carioca” type: G (6%), E (89%) and G×E (6%); and for the “Black” type: G (11%), E (68%), and G×E (21%). These results support our theory that a better understanding of E-related causes may support better breeding decisions and cultivar targeting, hypothesizing that this lack of environmental information might limit genetic progress in certain seasons and regions.

Despite the paradigm of G×E being a limiting factor for cultivar targeting, we demonstrate that climate prediction may improve the understanding of environmental limits and how future breeding research must overcome it. We envisage that some certain climate factors have drastically limited common beans yield, which may indirectly affect optimizing strategies for selecting the most adapted cultivars, such as (1) using envirotyping to characterize the environment of trials and verify whether it (2) defines the best stages, regions, and seasons for each market segment for phenotyping that are important eco-physiology traits, such as transpiration rate and biomass accumulation, which may also be used for more accurately training crop models capable of delivering a diverse set of solutions, such as ideotype design driving the selection and prediction of management conditions; and (3) taking advantage of indirect selection strategies using phenotypic records of different environments.

### 4.4 Fine-tunning environmental similarity among field trials, regions, and seasons

The modern plant breeding triangle (Crossa et al., 2021) takes advantage of genomics, enviromics, and phenomics in prediction-based selections, such as whole-genome prediction (WGP, Meuwissen *et al*., 2001), WGP with CGM (Crop Growth Model, Cooper et al., 2016; Messina et al., 2018), and enviromics prediction (Resende et al., 2021) by thematic maps of adaptation (Costa-Neto et al., 2020). At some point, each of those strategies uses two or three bases of the plant breeding triangle, previously described. Here, our results considered GAM approaches for understanding the climate limits of adaptation, which opens new pathways in common beans breeding research aiming to better explore the “triangle.” For example, GAM is a good performance model to identify how the response variable (grain yield) is modulated for a certain range of environmental covariables (EC), leading not only to demonstrating which ECs were the most important to explain GY, but also to a range of non-linearity that is a putative suggestion of the range of adaptation by a certain significant EC. This approach can be used for selecting the most important ECs in genomic-enabled reaction-norm studies involving WGP for predicting novel genotype-by-environment (G×E) conditions (Jarquín et al., 2014; Morais Júnior et al., 2018; Costa-Neto et al., 2021a). Observing our results, we hypothesize that different patterns of reaction-norm must be modeled for each region and season, perhaps ignoring grain type (it was not significant in most scenarios analyzed here). Another example relies in phenomics, in which the same GAM can be used to select wavelengths for predictive purposes.

We envisage that an interesting use of the climate prediction approach is to support the design of target envirotypes (environmental types) for environment assemblies (Costa Neto et al., 2021b). To derive structures of environmental relations among field trials, there is the concept of environment assembly inspired by the theory of “tolerance limits of adaptation,” also known as Shelford Law (Shelford, 1931). The Shelford Law preconizes that for some environmental gradient there are different adaptation zones according to genetic-oriented tolerance limits to a specific germplasm or species. There are death zones for a drastically lacking or for an excess of some environmental factor. Similarly, there are stress zones within which the limiting or exceeding factor compromises some biological process of growth and development (and consequently yield) but does not lead the crops to death.

The optimum conditions and the range of adaptation to key envirotypes can be achieved using the GAM-based strategy proposed here. Thus, the process of envirotyping (collecting, processing, and organizing data in an ecophysiologically smart way) undergoes from the definition of adaptation zones for a particular environmental factor (e.g., air temperature, solar radiation) to a particular germplasm in a particular region or season. In addition, a direct application of this information might rely on research transferability from different areas or seasons. For example, a research in the South region on some location and the some envirotype pattern and another research in the Midwest region can take advantage of similarities to accelerate field evaluations and selection of adapted cultivars.

## ACKNOWLEDGMENTS

AB Heinemann acknowledges support from “Fundação de Amparo à Pesquisa do Estado de Goiás” (FAPEG -PRONEM/FAPEG/CNPq) and “Conselho Nacional de Desenvolvimento Científico e Tecnológico” (CNPq –Edital Universal –Processo -408025/2018-2

## CONFLICT OF INTEREST

The authors declare that the research was conducted in the absence of any commercial or financial relationships that could be construed as a potential conflict of interest.

## CRediT AUTHOR STATEMENT

ABH and DHDM conceived the theory; ABH and IKF generated the dataset and performed the data analysis ; ABH and GCN wrote the manuscript, RFN and GCN revised the text, figures and tables, which all the authors finally edited.

## Notes

### Competing Interest Statement

The authors have declared no competing interest.

