## Supplementary for "Enviromic prediction is useful to define the limits of climate adaptation: A case study of common beans in Brazil"

**RUNTITLE**: Climate prediction of beans adaptation

**Highlights**

- We developed an ‘enviromic prediction’ approach based on Generalized Additive Models (GAM) for a large-scale environmental covariate data and grain yield
- We verified the ability of GAM-based models to explain the climate driver grain yield variation and performed accurate predictions for diverse production scenarios (four regions, three seasons, and two grain types)
- Air temperature (maximum and minimum), accumulated solar radiation, and rainfalls are mostly associated as the main drivers of GY variation in most regions.
- Climate influence in common beans adaptation is more evident during the vegetative for some seasons, while more impressive for reproductive stages for other seasons

**Table S1**. Variance components (G+E+GE) for “Carioca” and “Black” across regions collected from Rocha et al. (2020), and Pereira et al. (2017).

| Common bean Type | Region |  | Variance components | | | |
| --- | --- | --- | --- | --- | --- | --- |
| “Carioca” | Northeast | Source | MS | Df | SS | Importance SS |
|  |  | G | 240470 | 10 | 2404704 | 6% |
|  |  | E | 17964034 | 2 | 35928069 | 89% |
|  |  | GE | 113180 | 20 | 2263613 | 6% |
|  |  | TOTAL (G+E+GE) = | | | 405963876 |  |
|  |  | Source | MS | Df | SS | Importance SS |
| “Black” | Northeast | G | 344053 | 14 | 4816749 | 11% |
|  |  | E | 14402964 | 2 | 28805929 | 68% |
|  |  | GE | 310201 | 28 | 8685631 | 21% |
|  |  | TOTAL (G+E+GE) = | | | 42308310 |  |
|  |  | Source | MS | df | SS | Importance SS |
| “Carioca” | Southeast | G | 2514763 | 15 | 37721445 | 2% |
|  |  | E | 40317209 | 35 | 1411102315 | 88% |
|  |  | GE | 437526 | 362 | 158384412 | 10% |
|  |  | TOTAL (G+E+GE) = | | | 1607208172 |  |
|  |  | Source | MS | df | SS | Importance SS |
| “Carioca” | Midwest | G | 1306778 | 15 | 19601670 | 2% |
|  |  | E | 24185199 | 36 | 8.71E+08 | 82% |
|  |  | GE | 513422 | 332 | 1.7E+08 | 16% |
|  |  | TOTAL (G+E+GE) = | | | 1.06E+09 |  |

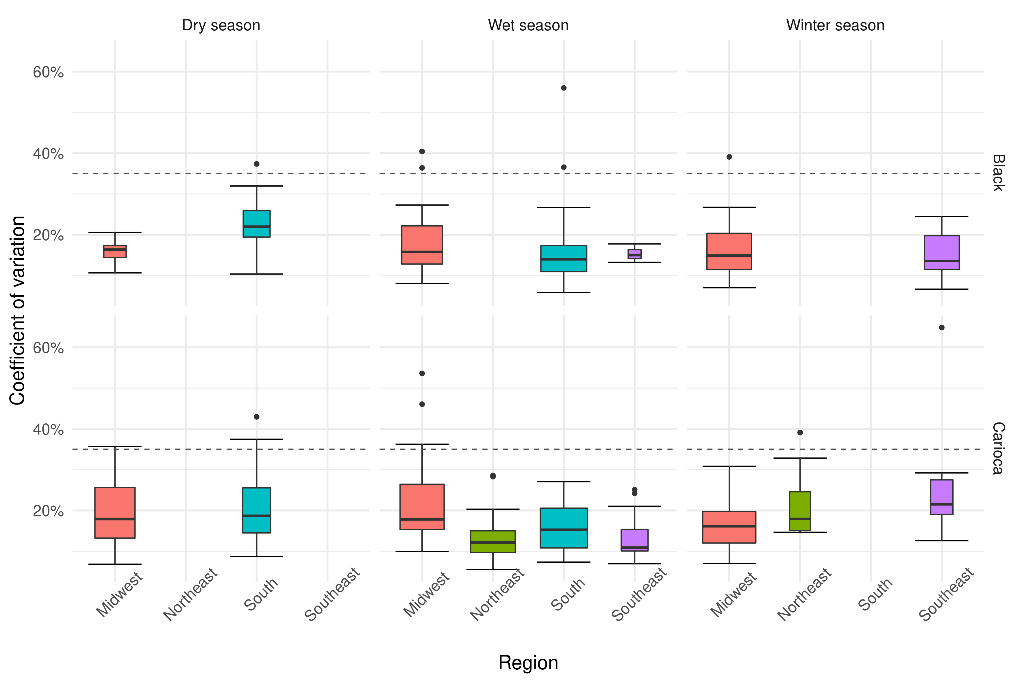

**Figure S1**. Coefficient of variation (%) for all trials used in this study across seasons (top panel), common bean types (right panel), and production regions. The dashed line shows the limit of coefficient of variation for 35%. At each boxplot, the values of the boxes represent the 1st quartile (25%, bottom limit), second quartile (50% or median, bold horizontal line), and third quartile (75%, top limit). The bars indicate the minimum and maximum observed values for each distribution of coefficients of variation values across the different setups.

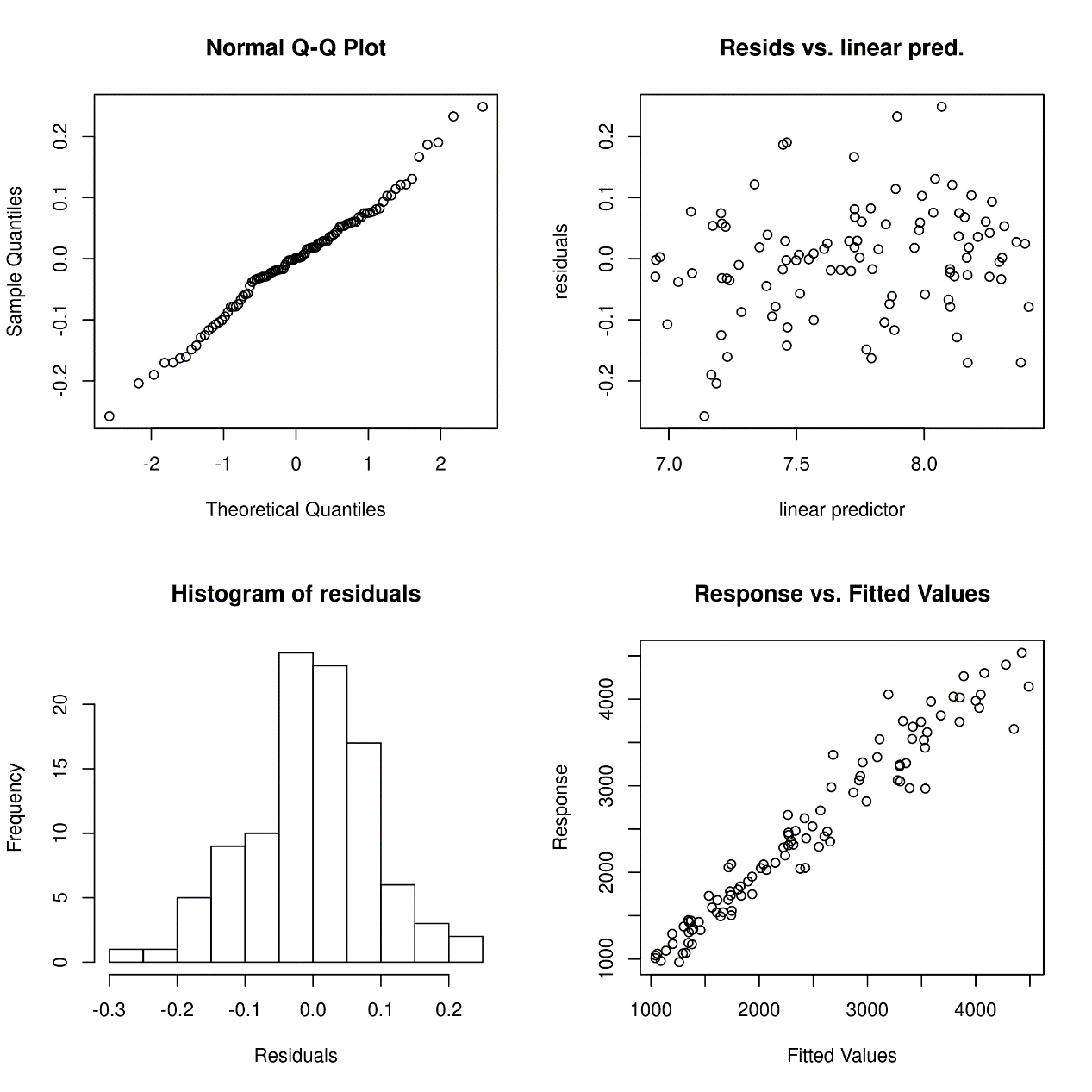

**Figure S2.** GAM performance for the south region.

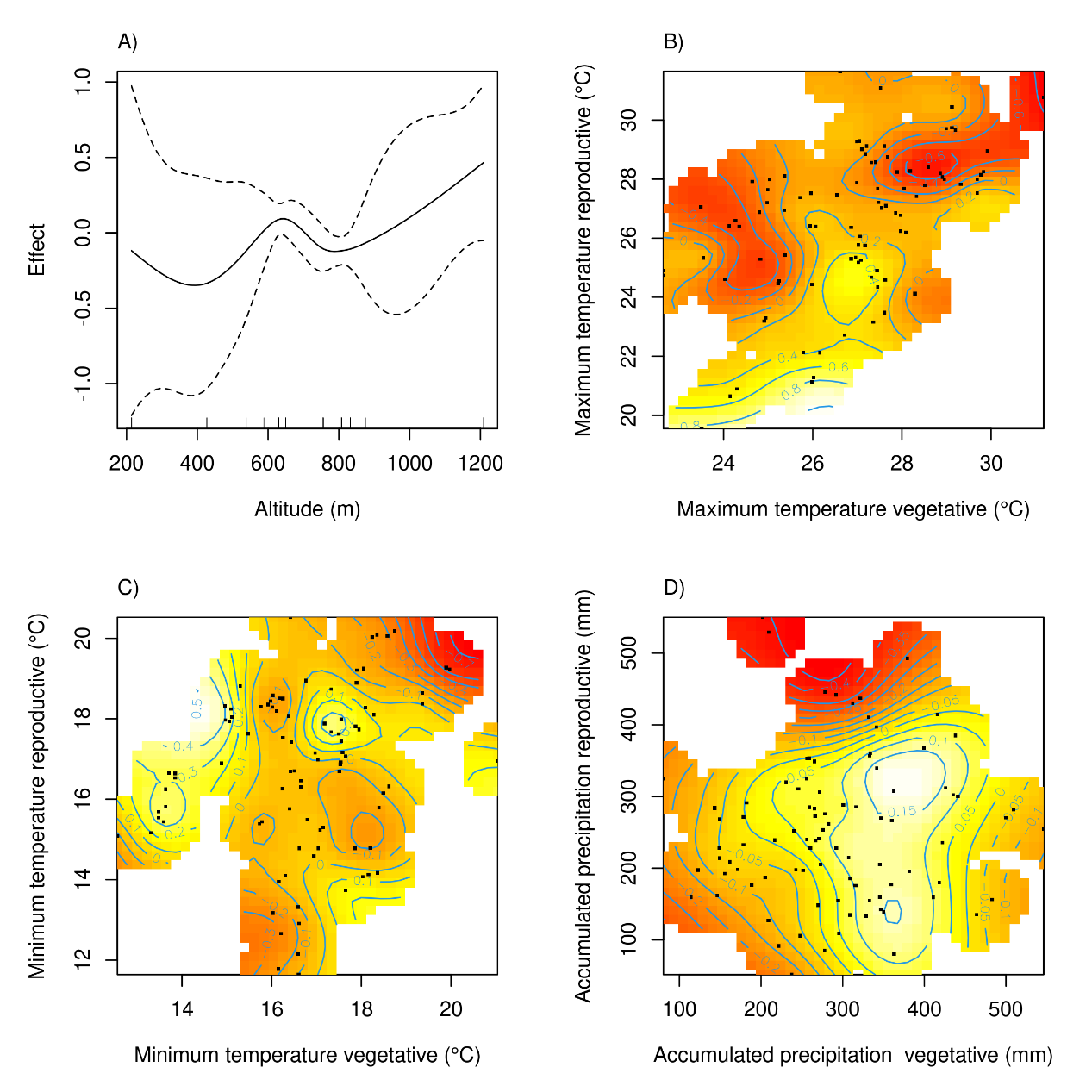

**Figure S3**. Smooth terms for South region considering all seasons (dry and wet): A) Altitude (m), B) maximum and C) the minimum temperature at vegetative and reproductive phases, and D) precipitation accumulated at vegetative and reproductive phases. Black points represent the observed common beans yield from trials; blue lines show the contours lines, numbers in contours lines represent the effect on yield as well as the colors, yellow positive and red negative effect.

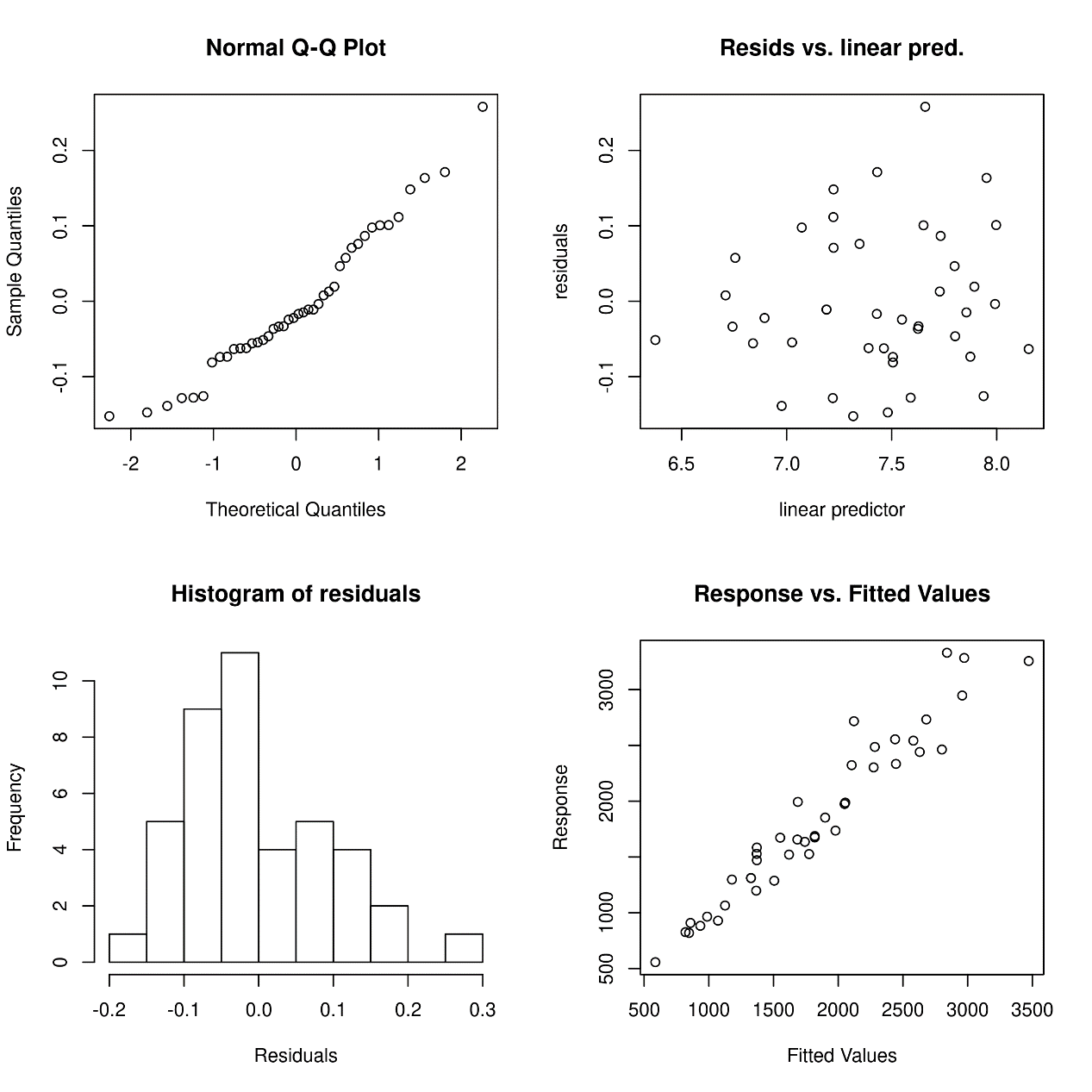

**Figure S4.** GAM performance for the southeast region.

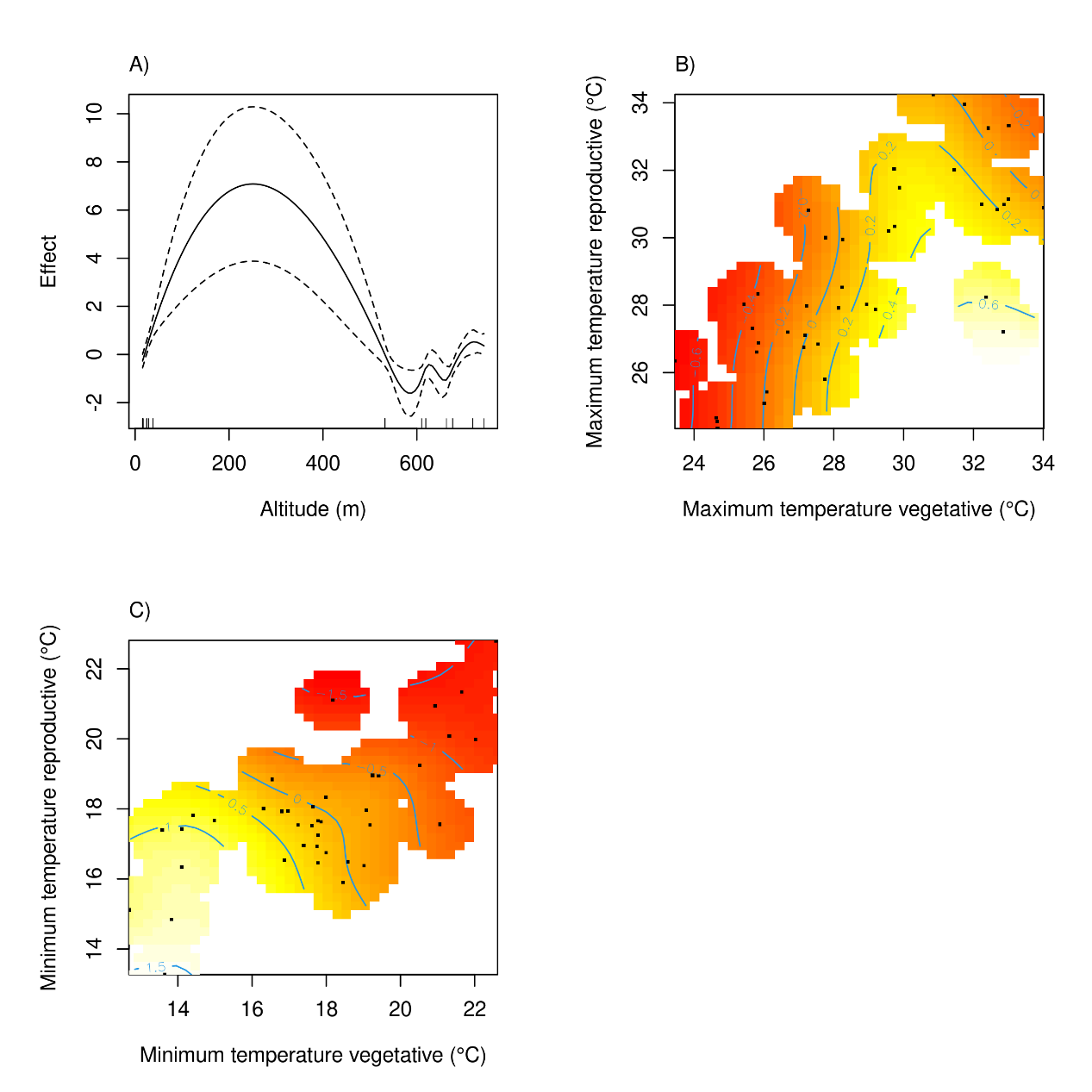

**Figure S5.** Smooth terms for Southeast region considering all seasons (wet and winter) A) Altitude (m), B) maximum and C) minimum temperature for vegetative and reproductive phases, for Southeast region. Black points represent the observed common beans yield from trials; blue lines show the contours lines, numbers in contours lines represent the effect on yield as well as the colors, yellow positive, and red negative effect.

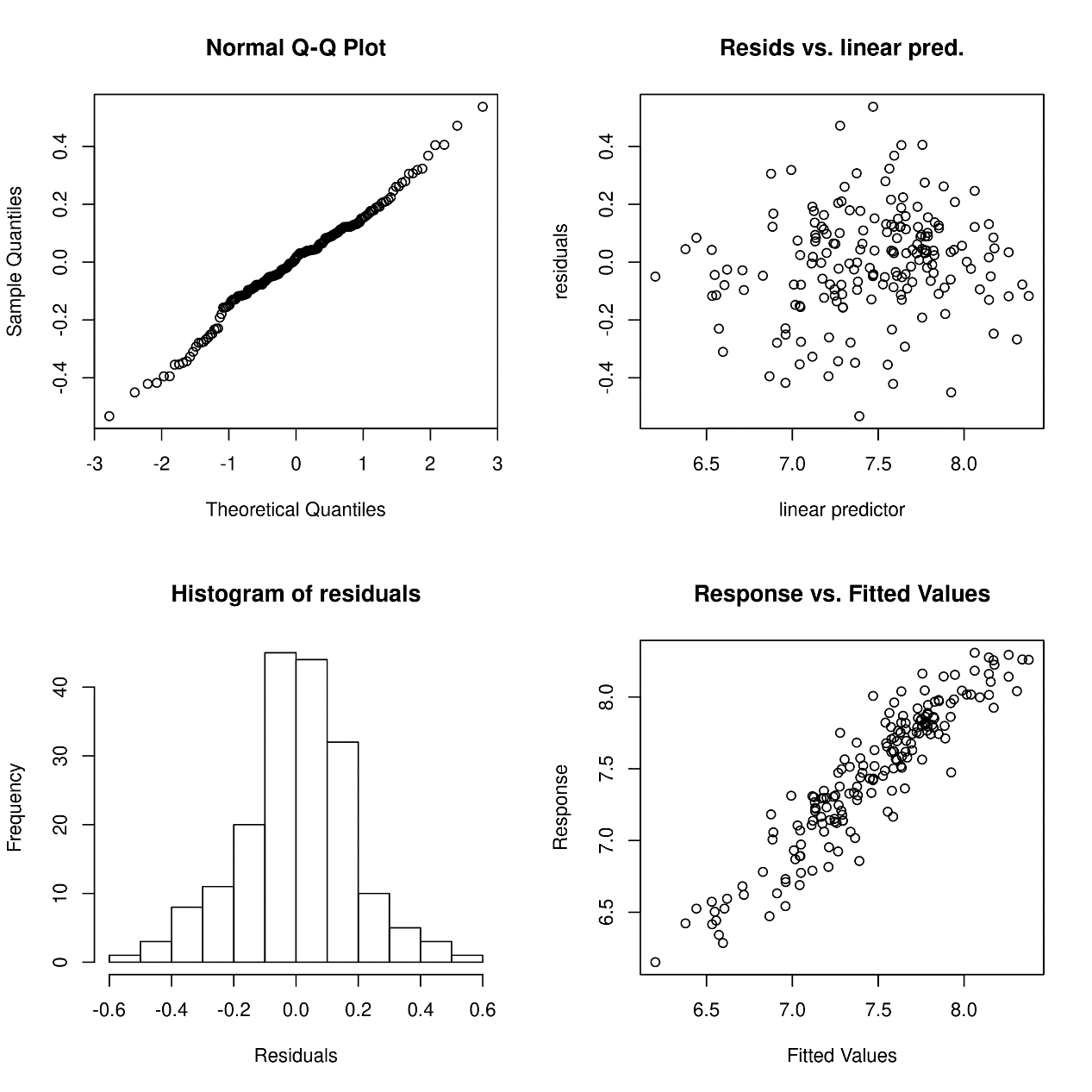

**Figure S6.** GAM performance for Midwest region considering all seasons (wet, dry, and winter).

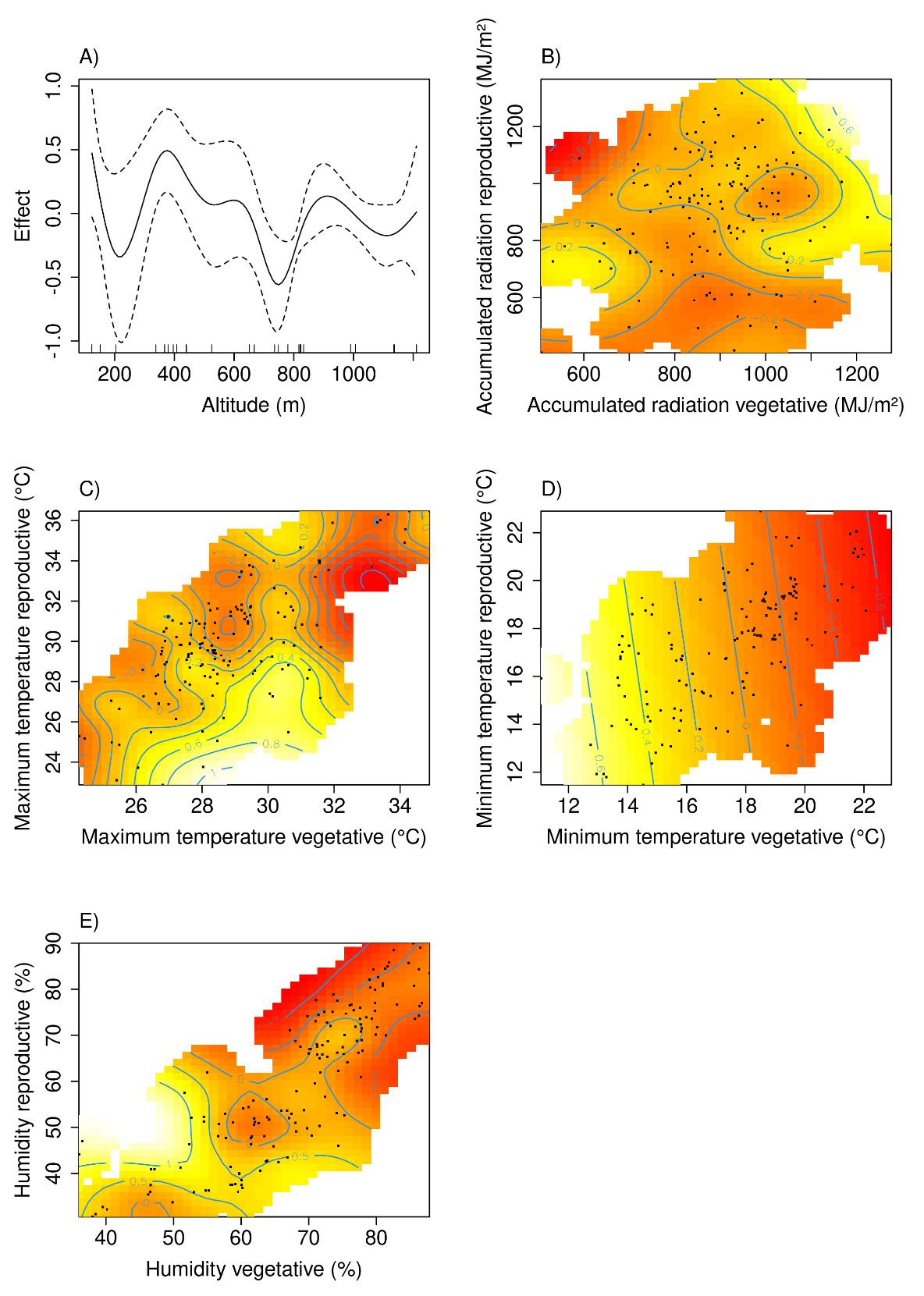

**Figure S7.** Smooth terms for Midwest region considering all seasons (wet, dry, and winter), A) Altitude (m), B) solar radiation accumulated, C) maximum and D) minimum temperature for vegetative and reproductive phases, E) relative humidity for vegetative and reproductive phases for vegetative and reproductive phases. Black points represent the observed common beans yield from trials; blue lines show the contours lines, numbers in contours lines represent the effect on yield as well as the colors, yellow positive, and red negative effect.

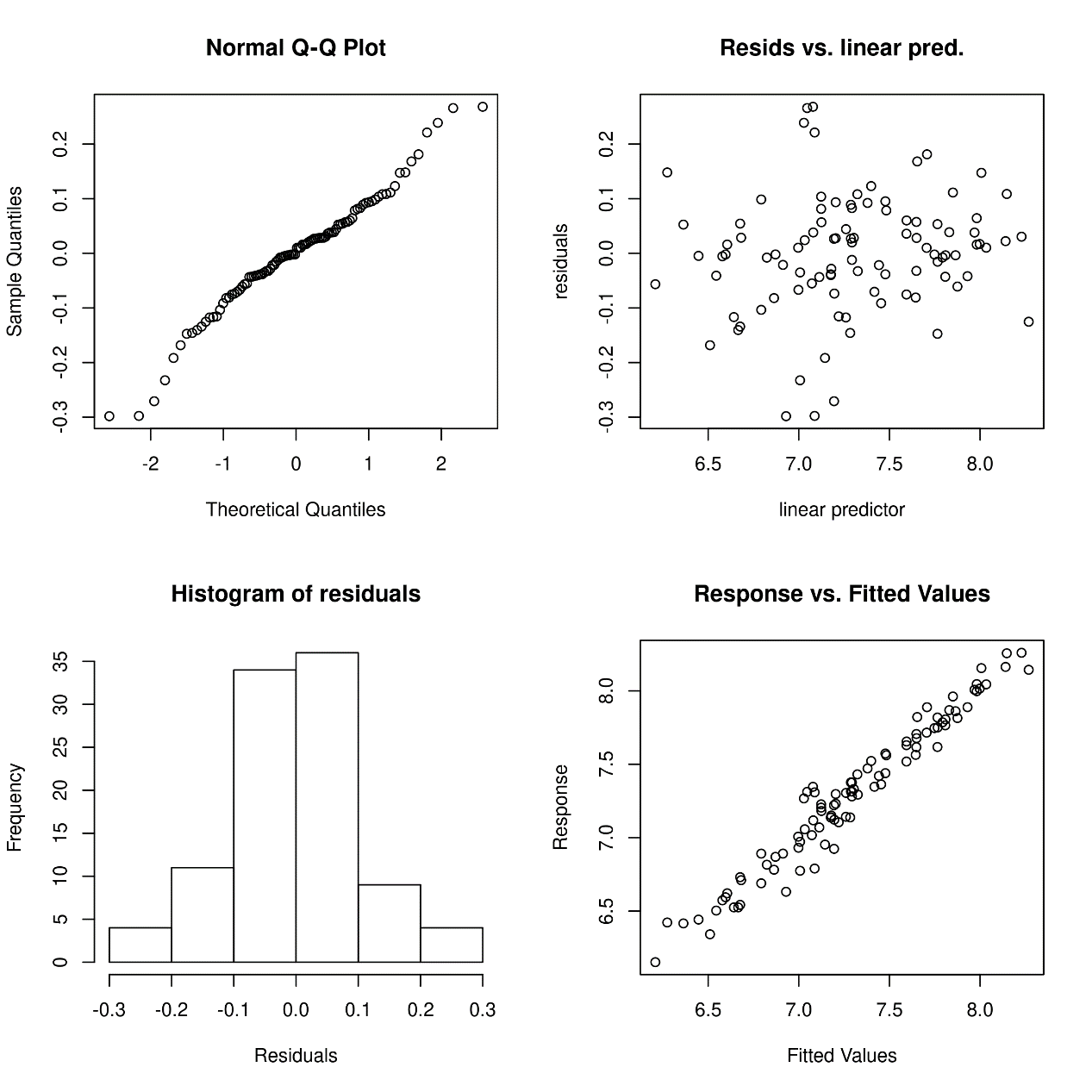

**Figure S8.** GAM performance for Midwest region considering only wet and dry seasons (rainfed seasons).

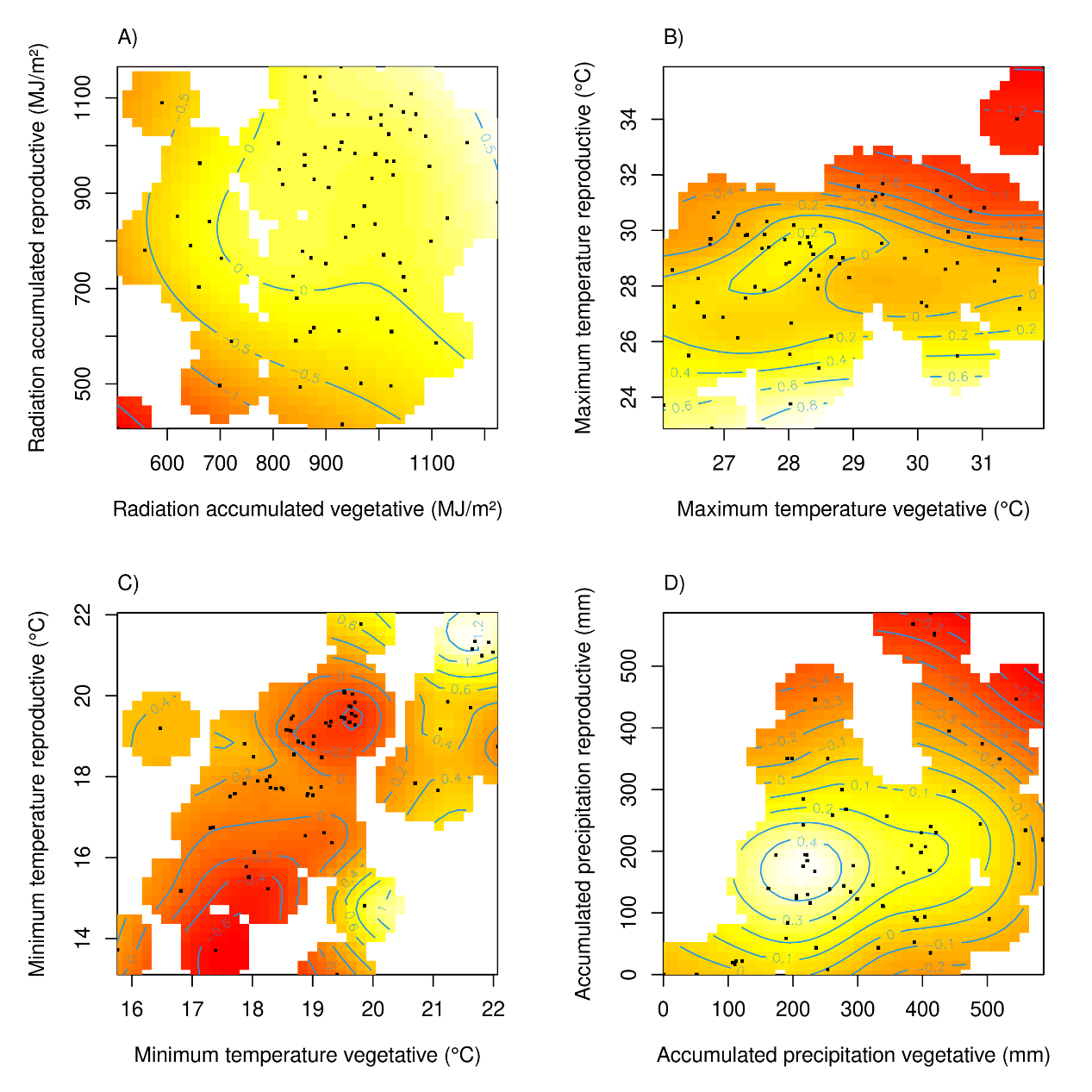

**Figure S9**. Smooth terms for Midwest region considering only wet and dry seasons (rainfed seasons): A) Altitude (m), B) maximum and C) minimum temperature for vegetative and reproductive phases, and D) accumulated precipitation for vegetative and reproductive phases. Black points represent the observed common beans yield from trials; blue lines show the contours lines, numbers in contours lines represent the effect on yield as well as the colors, yellow positive and red negative effect.

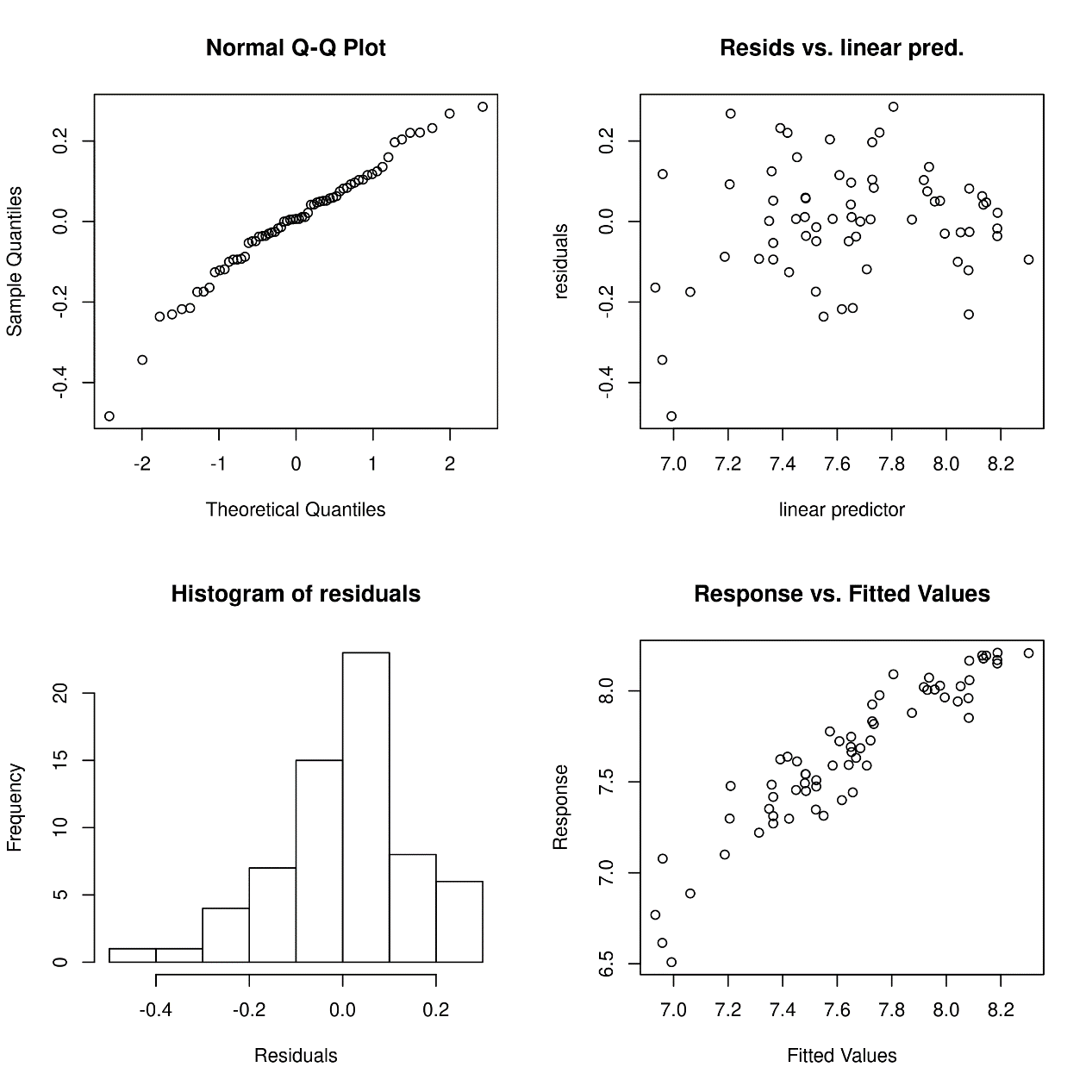

**Figure S10.** GAM performance for Northeast region considering all seasons (wet and winter).

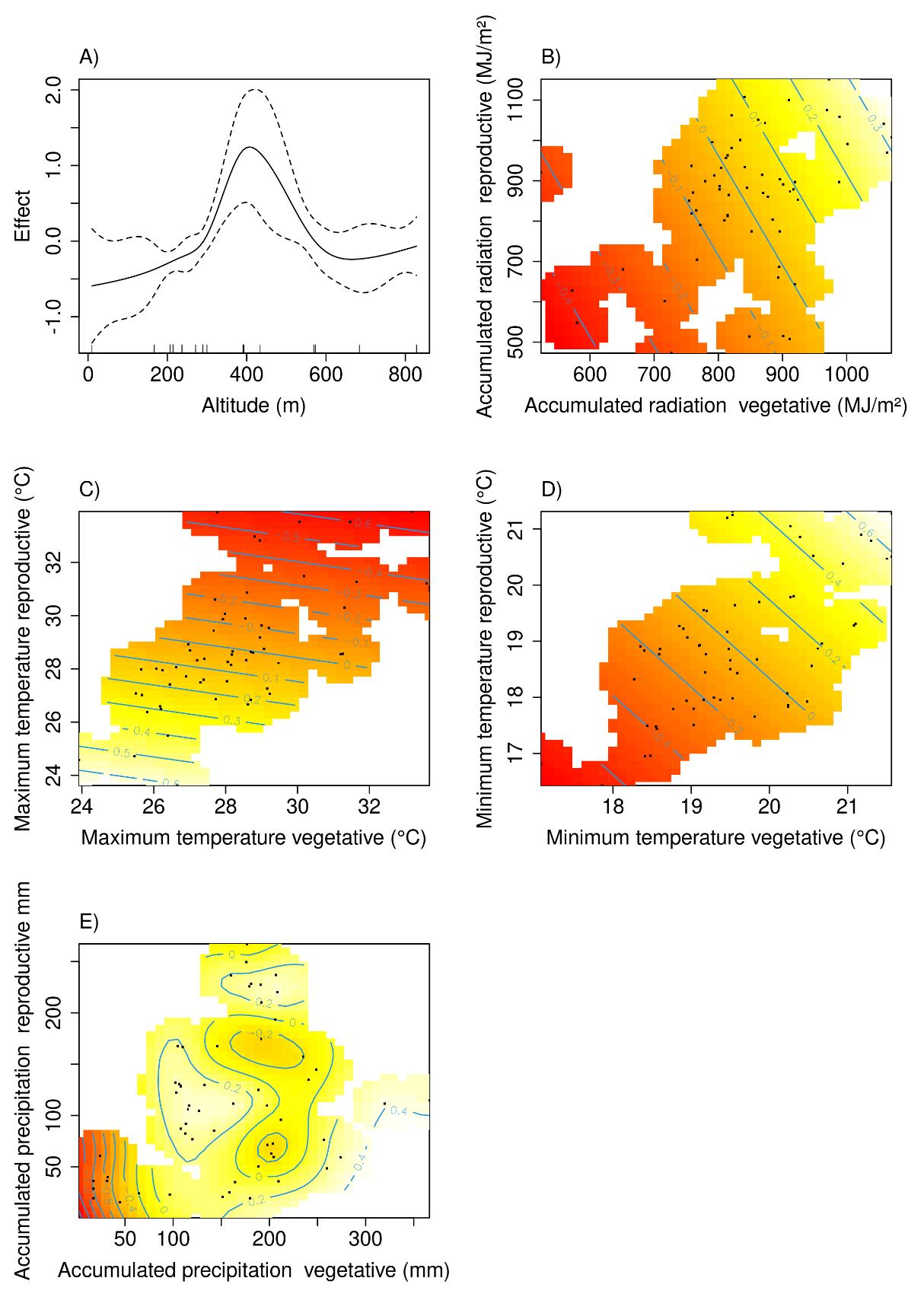

**Figure S11.** Smooth terms for Northeast region considering all seasons (wet and winter): A) Altitude (m), B) accumulated solar radiation D) maximum and E) minimum temperature for vegetative and reproductive phases, and F) accumulated precipitation for vegetative and reproductive phases. Black points represent the observed common beans yield from trials; blue lines show the contours lines, numbers in contours lines represent the effect on yield as well as the colors, yellow positive and red negative effect.
